# Viral and host network analysis of the human cytomegalovirus transcriptome in latency

**DOI:** 10.1101/2024.05.21.594597

**Authors:** Donna Collins-McMillen, Diogo De Oliveira Pessoa, Kristen Zarrella, Christopher J. Parkins, Michael Daily, Nathaniel J. Moorman, Jeremy P. Kamil, Patrizia Caposio, Megha Padi, Felicia D. Goodrum

**Affiliations:** BIO5 Institute, University of Arizona, Tucson, Arizona, United States of America; Department of Immunobiology, University of Arizona, Tucson, Arizona, United States of America; Bioinformatics Shared Resource, Arizona Cancer Center, University of Arizona, Tucson, Arizona, United States of America; Department of Microbiology and Immunology, University of North Carolina at Chapel Hill, Chapel Hill, North Carolina, United States of America; Department of Microbiology and Immunology, Louisiana State University Health Sciences Center, Shreveport, Louisiana, United States of America; Vaccine and Gene Therapy Institute, Oregon Health Science University, Beaverton, Oregon, United States of America; University of Arizona Cancer Center, University of Arizona, Tucson, Arizona, United States of America; Department of Molecular and Cellular Biology, University of Arizona, Tucson, Arizona, United States of America

## Abstract

HCMV genes *UL135* and *UL138* play opposing roles regulating latency and reactivation in CD34^+^ human progenitor cells (HPCs). Using the THP-1 cell line model for latency and reactivation, we designed an RNA sequencing study to compare the transcriptional profile of HCMV infection in the presence and absence of these genes. The loss of *UL138* results in elevated levels of viral gene expression and increased differentiation of cell populations that support HCMV gene expression and genome synthesis. The loss of *UL135* results in diminished viral gene expression during an initial burst that occurs as latency is established and no expression of eleven viral genes from the UL*b*’ region even following stimulation for differentiation and reactivation. Transcriptional network analysis revealed host transcription factors with potential to regulate the UL*b*’ genes in coordination with pUL135. These results reveal roles for *UL135* and *UL138* in regulation of viral gene expression and potentially hematopoietic differentiation.

## Introduction

Human cytomegalovirus (HCMV) establishes a lifelong persistent infection that is marked by periods of latency and reactivation. Viral gene expression is limited during latency, allowing the virus to escape immune clearance and persist in the host (*1*). In the healthy host, reactivation occurs sporadically and involves production and shedding of progeny virus with little to no pathogenesis. In contrast, HCMV infection or reactivation in the immunocompromised or immune naive individual can lead to severe morbidity and mortality (*2*).

The HCMV genome remains incompletely annotated. With its ∼236 kilobase pair double-stranded DNA genome, HCMV is the largest of all viruses known to infect humans and has been shown to encode over 170 viral proteins (*1*). The introduction of HCMV gene arrays and next-generation sequencing technologies such as RNA-Seq has revolutionized our ability to study HCMV gene expression at a genome-wide level during both lytic (*3–7*) and latent infections (*8–14*). A 2011 transcriptome (*4*) and a 2012 ribosomal profiling study (*15*) revealed the enormous and complex coding capacity of HCMV with an estimated potential for 751 open reading frames (ORFs) in total. Many of these ORFs were reported to contain splice junctions or alternative transcript start sites that add further complexity to the viral gene expression program (*4, 15*).

For example, alternative splicing that gives rise to distinct gene products or isoforms of a single gene has been characterized in a small number of HCMV genes, mostly from the immediate early gene regions (*4, 16, 17*). Additional complexity arises through usage of alternative initiation or termination signals to produce multiple peptides from a single transcriptional unit (*17*). These transcripts can be polycistronic, encoding multiple genes from a single transcript, or monocistronic, encoding multiple isoforms of a single gene (*17*). In the case of *UL4* (*17–19*), *UL44* (*17, 20*), and *UL122-UL123* (*21*), multiple transcripts are made that encode identical gene product(s) but have distinct 5’ untranslated regions. This mechanism allows for the regulation of gene expression by alternative promoters under different contexts of infection. *UL4*, *UL44* and *UL122-UL123* are each subjected to a degree of temporal control, as different combinations of the promoters are active during immediate early, early, and late phases of lytic infection (*18–21*). More recently, we showed that alternative promoters in intron A of the *UL122-UL123* locus are active in hematopoietic cells during latency, whereas activity of the canonical major immediate early promoter (MIEP) remains low (*22, 23*). The alternative promoters control the accumulation of the critical MIE transactivators during reactivation from latency in our THP-1 and CD34^+^ HPC models (*22, 23*); however this appears to be dependent on cell type and reactivation stimulus (*24, 25*). These findings suggest that the virus has evolved a sophisticated strategy for initiating the replicative cycle in response to different cellular stimuli or contexts.

The UL*b*’ region of the HCMV genome is another hotspot for complex regulation of gene expression. UL*b*’ contains four polycistronic loci that each produce sets of 3’ co-terminal transcripts encoding multiple combinations of viral genes during different temporal phases of infection (*17*). The *UL133-UL138* locus that modulates latency and reactivation is among these and has a high degree of complexity in expression of its gene products. Transcripts expressed with early kinetics encode some combination of all four gene products, pUL133, pUL135, pUL136, and pUL138 (*26*). During the late phase of infection, transcripts encoding only two of these proteins, pUL136 and pUL138, are produced and pUL136 accumulates to maximal levels only following commitment to vDNA synthesis (*26*). This level of temporal control hints at a complex and highly responsive regulation of these genes to meticulously govern the switch between latent and replicative infection in response to cellular cues.

We have previously reported that the HCMV genes *UL135* and *UL138* are expressed with early kinetics and their gene products play opposing roles in regulating latency and reactivation in primary CD34^+^ human progenitor cells (HPCs) (*27–30*). An HCMV recombinant virus that fails to express *UL135* is defective for reactivation from latency and replication in CD34^+^ HPCs, whereas a recombinant that fails to express *UL138* is defective for establishment of latency and actively replicates in CD34^+^ HPCs in the absence of a stimulus for reactivation. Here, we used these recombinant viruses to conduct a large-scale RNA sequencing (RNA-Seq) study to better define the roles of *UL135* and *UL138* in infection of hematopoietic cells.

For these studies, we used the monocyte-like THP-1 cell line to model a latent infection. THP-1 cells have been an important tool for exploring the regulation of viral gene expression and genome silencing in HCMV latency (*22, 23, 31–39*) because they allow for synchronous expression, silencing, and re-expression of viral genes given that they lack the heterogeneity of primary cell models, which has been a major issue in defining the patterns of gene expression associated with latency. In comparing the distinct transcriptional profiles associated with the expression of *UL135* and *UL138* in infection, we aim to further define the roles of *UL135* and *UL138* in viral infection in hematopoietic cells and define key viral and host factors that coordinate the switch between latent and replicative states.

This transcriptome defines key aspects of viral gene expression during HCMV infection in THP-1 cells. A burst of viral gene expression is observed following infection and is silenced within days for the establishment of the latent-like infection. However, in cells infected with a *UL135*-mutant virus, the initial burst of gene expression does not occur. Re-expression of viral genes is uniformly induced by TPA; however, the *UL135*-mutant virus fails to express a block of eleven UL*b*’ genes in THP-1 cells, although they are readily detected during *UL135*-mutant virus infection of replication-permissive fibroblasts. Transcriptional network analysis revealed host transcription factors that are modulated by pUL135 and predicted to regulate the eleven UL*b*’ genes driven by pUL135. These transcription factors may drive reactivation from latency and, therefore, represent promising cellular targets for controlling viral reactivation in a clinical setting. In the case of the *UL138*-mutant virus infection, gene expression is overall increased relative to the wild-type virus, although the pattern of gene expression is similar. Importantly, *UL138*-mutant virus infection resulted in increased spontaneous differentiation of THP-1 cells into an adherent cell that expressed higher levels of viral transcripts compared to cells that remained in suspension. These results define roles for *UL135* and *UL138* in the regulation of viral gene expression and possibly hematopoietic differentiation.

This work furthers our understanding of the roles *UL138* and *UL135* play in regulating HCMV infection and replication in hematopoietic cells. In addition, we have used this data set to identify host factors potentially underlying the changes in viral gene expression for future work. This transcriptome will be an important resource going forward for guiding exploration of the latent versus replicative HCMV transcriptional programs and the underlying virus/host interactions that modulate the complex viral persistence strategy.

## Results

### Analysis of the *UL135*- and *UL138*-dependent control of the HCMV transcriptome

We previously engineered recombinant viruses containing 5’ stop codon substitutions in the *UL135* (Δ*UL135*_STOP_) and *UL138* (Δ*UL138*_STOP_) genes in the HCMV TB40/E strain to disrupt synthesis of their corresponding proteins (*29*). Δ*UL135*_STOP_ fails to reactivate from latency, as it produces lower levels of productive virus from latently infected CD34^+^ HPCs following a reactivation stimulus in comparison to WT infection (Figure 1A). In contrast, the Δ*UL138*_STOP_ recombinant fails to establish latency and produces greater levels of viral progeny when compared to WT infection both prior to and following reactivation stimulus. We have further shown that *UL135* and *UL138* have an antagonistic relationship and that *UL138* expression restricts HCMV reactivation in the absence of *UL135* (*29*), which is due at least in part to the opposing regulation of EGFR turnover and signaling (*30*). The global changes in patterns of viral gene expression associated with these phenotypes are unknown. Therefore, we analyzed the HCMV transcriptome over a time course in THP-1 cells infected with wildtype (WT), Δ*UL135*_STOP_ or Δ*UL138*_STOP_ compared to mock-infected cells. Cells infected in their undifferentiated state were cultured for 5 days to establish a latent infection. On day 5, infected cell cultures were divided; half were treated with the vehicle DMSO (undifferentiated) and half were treated with phorbol ester TPA (differentiated) to promote monocyte-to-macrophage differentiation and trigger re-expression of viral genes. Total RNA was sequenced at key time points for four biological replicates.

**Figure 1.**
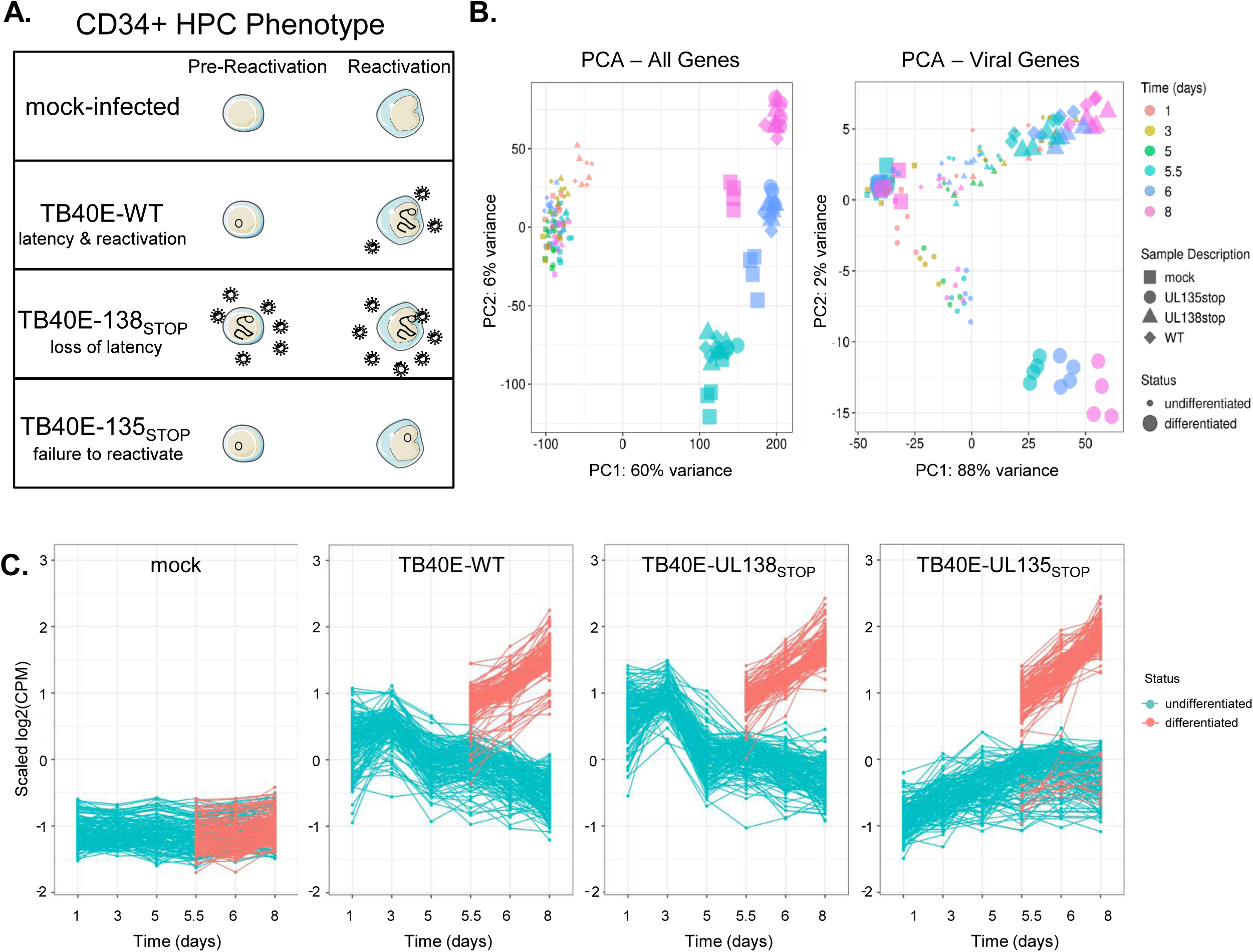
Analysis of the *UL135*- and *UL138*-dependent control of the HCMV transcriptome. **A)** A depiction of the samples included in this analysis and the phenotype of each virus in our CD34^+^ HPC model. Mock-infected cells were used to establish a baseline for regulation of cellular genes. Wildtype (WT) virus establishes latency in CD34^+^ HPCs and reactivates in response to cytokine stimulus. The Δ*UL138*_STOP_ recombinant is replicative both prior to and following reactivation stimulus (loss of latency) and the Δ*UL135*_STOP_ recombinant has low replication both prior to and following reactivation stimulus (failure to reactivate). **B)** Principal Component Analysis (PCA) plots were made using the ggplot2 package (*60*). Plots were made for both cellular and viral genes (left) and for viral genes only (right). Treatment groups include mock-infected as well as samples infected with WT, Δ*UL135*_STOP_, or Δ*UL138*_STOP_ HCMV. Each treatment group consists of samples collected at 1, 3, and 5 days post infection (dpi) and at 5.5, 6, and 8 dpi treated with either DMSO control or TPA to induce cellular differentiation and viral reactivation. **C)** A time course of viral gene expression was made for each treatment group. Each set of data points connected by a single line represents one HCMV gene. Data were scaled to log2 counts per million (CPM) as a function of gene count.

The resulting data were first reduced to two dimensions by Principal Component Analysis (PCA) (Figure 1B) to identify which factors produce the greatest variance across the data set. When both cellular and viral genes are included in the analysis (Figure 1B, PCA-All Genes), 60% of total variance is due to differentiation status of the cells. The undifferentiated samples largely cluster together, with a small outlier effect for WT-infected and Δ*UL138*_STOP_-infected at 1 day post infection (dpi). Among the differentiated samples, most of the separation along the y-axis, representing 6% of total variance, is attributed to time point (5.5, 6, or 8 dpi), reflective of changes in gene expression that occur as the cells further differentiate. Differences are also seen between mock-infected cells and those infected with any of the three viruses. It is unsurprising that the greatest source of variance in the complete transcriptome is differentiation status, given the outsized contribution of cellular genes compared to viral genes (particularly in latency-associated cells where viral gene expression is restricted) and the number of cellular gene expression changes that occur during monocyte-to-macrophage differentiation.

To better understand variance driven by the viral transcriptome, we assembled a second PCA of viral genes alone (Figure 1B, PCA – Viral Genes). Again, the greatest variance (PC1, 88%) is associated with differentiation state. This effect is more pronounced in the Δ*UL135*_STOP_-infected samples, whereas the WT-infected and Δ*UL138*_STOP_-infected samples have more overlap between the differentiated samples at all time points and the undifferentiated samples at 1 and 3 dpi as latency is being established. The Δ*UL135*_STOP_-infected samples cluster separately from the WT-infected and Δ*UL138*_STOP_-infected samples along the y-axis, which accounts for 2% of total variance across the data set. These results suggest that there are important differences in the viral gene expression program of the Δ*UL135*_STOP_ recombinant virus when compared to the WT parental virus or the Δ*UL138*_STOP_ recombinant.

We next plotted the viral read counts for each experimental condition over the time course of infection to discern differences in viral gene expression patterns (Figure 1C). Each series of plotted points connected by a line represents a single viral gene. Values were normalized to average read count for the same gene across the data set and log-transformed so that each increment of 1 on the y-axis represents a two-fold change in viral gene expression. As expected, read counts for viral genes in the mock-infected samples are low/undetected across the data set. Read counts in the WT-infected samples reveal an initial burst of viral gene expression at 1 dpi that decreases as latency is established (undifferentiated) and the re-initiation of viral gene expression following TPA treatment (differentiated). Viral gene expression follows a similar pattern in the Δ*UL138*_STOP_-infected samples; however, Δ*UL138*_STOP_ read counts are higher than WT read counts at each time point. We were surprised that the increase in viral gene expression for Δ*UL138*_STOP_ was not greater, as our work in primary CD34^+^ HPCs demonstrated a clear increase in virus replication and a defect in establishing latency. This phenotype is further explored in a later section.

The most striking differences in viral gene expression are seen in the Δ*UL135*_STOP_-infected samples. Viral gene expression is lower from 1 to 5 dpi when latency is being established. Following a reactivation stimulus, viral gene expression increases similar to WT and Δ*UL138*_STOP_ infection, with the exception of a small group of viral genes that remain silenced.

Because the Δ*UL135*_STOP_ recombinant has been characterized as deficient for reactivation from latency in CD34^+^ HPCs, we postulated that this select group of viral genes might play an integral role in driving viral reactivation from latency. Intriguingly, these results suggest that the failure of Δ*UL135*_STOP_ to reactivate may not be due to a global failure to re-express viral genes, and instead hinges on the timely expression of a few key viral genes. These data further indicate a potential function for the initial burst of viral gene expression in the establishment of a latent infection that can later be reactivated.

### Viral genes cluster into distinct patterns of regulation during latency and reactivation

To analyze the expression kinetics of individual viral genes, we performed k-means clustering of viral reads detected across the data set (Figure 2A). Data were scaled to ensure that genes cluster together not solely because they are expressed at similar levels on average, but rather because they share similar expression dynamics, providing hints at co-regulation by common factors. Following the WT infection, viral genes are expressed at days 1 and 3 during the establishment of latency, then read counts decrease in subsequent undifferentiated samples and remain low following DMSO control treatment. In the differentiated samples, viral genes are re-expressed following TPA treatment, which triggers monocyte-to-macrophage differentiation. The genes in clusters 1 and 3 follow the expression pattern conventionally associated with latency and reactivation, where gene expression is comparatively low in the undifferentiated samples once latency is established and then increases following the reactivation stimulus. Genes that belong in cluster 2 are expressed at lower levels in the WT infection when compared to either of the recombinant viruses during both the maintenance of latency and following reactivation stimulus. Finally, the cluster 4 genes are dependent on *UL135* for their expression at any time point after infection, and particularly following TPA treatment.

**Figure 2.**
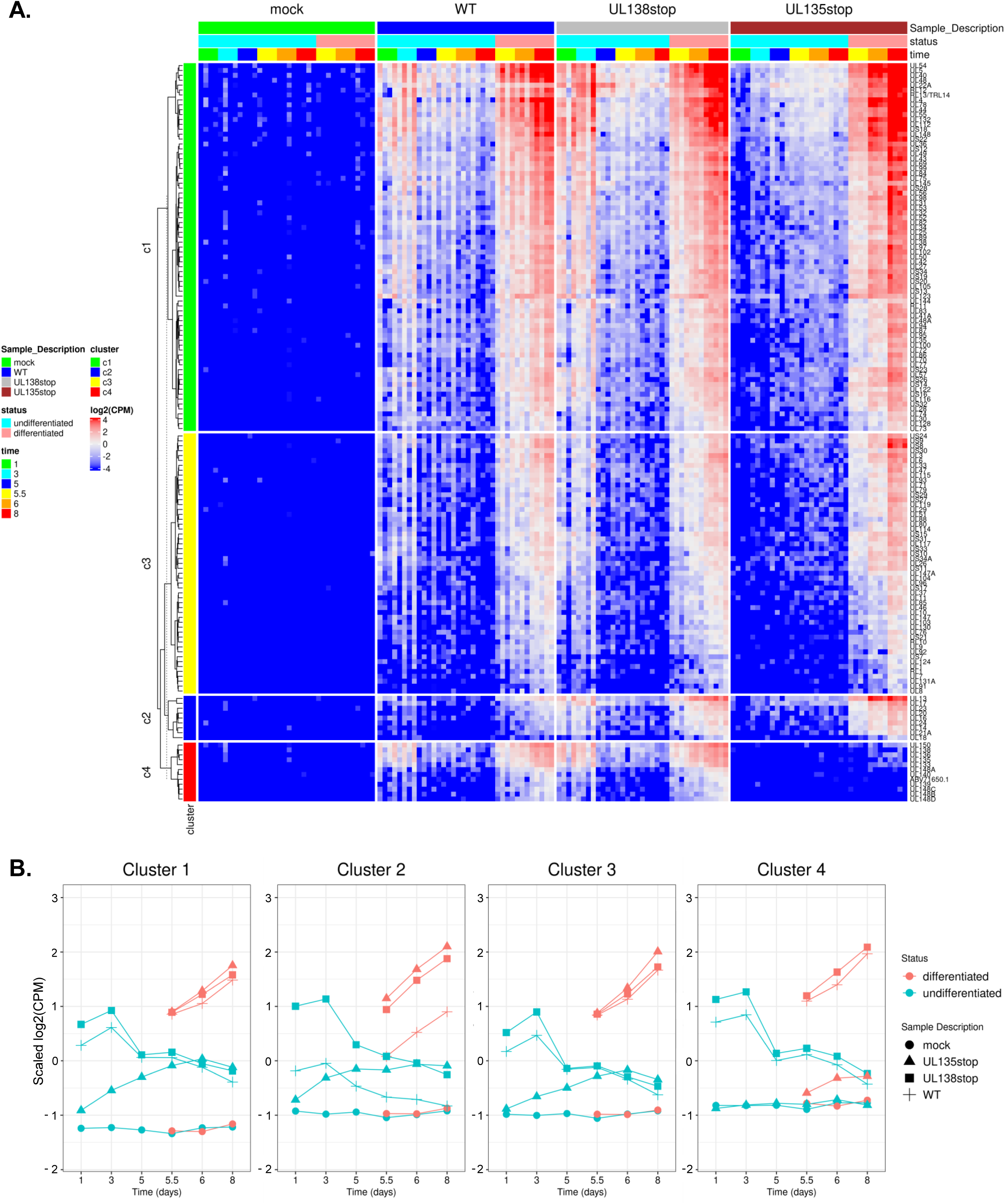
Viral genes cluster into distinct patterns of regulation during latency and reactivation. **A)** Clustering analysis was performed using the k-means approach and viral gene expression was visualized via heatmap with the *ComplexHeatmap* package from R (*64, 65*). Data were scaled to log2 CPM as a function of gene count. **B)** Average viral gene expression for each treatment group is shown by viral gene cluster.

To better appreciate cluster-specific differences in viral gene expression among the different viruses, read counts were averaged for each cluster of genes over the time course and plotted by individual infection group (Figure 2B). WT infection is characterized by a global initial burst of gene expression across all clusters at 1 dpi, which is silenced beginning after 3 dpi, and re-expressed following treatment with TPA. With respect to Δ*UL138*_STOP_, viral gene expression is increased relative to WT-infection, particularly at 1 and 3 dpi in all clusters. In addition, Δ*UL135*_STOP_ fails to express viral genes in all clusters at 1 and 3 dpi and also during the re-expression of cluster 4 genes following TPA treatment. Additionally, these plots clearly illustrate that cluster 2 gene expression following TPA is uniformly upregulated in both Δ*UL135*_STOP_ and Δ*UL138*_STOP_ infection relative to WT infection.

The gene expression dynamics defined by the transcriptome were confirmed by RT-qPCR of WT-infected THP-1 cells for a subset of viral genes. Viral transcripts were quantified at 1 and 7 dpi in each cluster, representing multiple kinetic classes (Figure S1). The expression of *UL123* and *UL86* (cluster 1), *UL17* (cluster 2), *US27* (cluster 3), and *UL135* and *UL138* (cluster 4) is consistent with the normalized expression shown in the clustering analysis (Figure 2).

### The Δ*UL138*_STOP_ loss of latency phenotype is pronounced in a subset of HCMV-infected hematopoietic cells

We were struck by the similarity between the WT and Δ*UL138*_STOP_ viral transcriptomes, given the replicative phenotype in CD34^+^ HPCs where the Δ*UL138*_STOP_ recombinant produces similar numbers of infectious progeny both prior to and following reactivation stimulus, demonstrative of a loss of latency (*27, 29, 30, 40*). To further explore this, we analyzed Δ*UL138*_STOP_ infection *in vivo* using humanized mice (Figure 3A). NOD-*scid* IL2Rγ_c_^null^ (huNSG) mice were sub-lethally irradiated and engrafted with human CD34^+^ HPCs prior to intraperitoneal injection of fibroblasts infected with *UL138*myc (parental virus expressing a myc-tagged variant of pUL138) or Δ*UL138*_STOP_. Stem cell mobilization and virus reactivation and dissemination was stimulated at 4 weeks in 5 of 10 mice using granulocyte colony stimulating factor (G-CSF). After 7 days, viral genomes were detected by qPCR in liver and spleen tissues. Untreated mice infected with *UL138*myc had low levels of HCMV genome detected in the spleen and liver, consistent with a latent infection, and viral genomes were increased in both tissues following G-CSF treatment, consistent with increased mobilization and reactivation from latency. In contrast, mice infected with the Δ*UL138*_STOP_ recombinant had similarly high HCMV genome copies in the liver and spleen both prior to and following G-CSF mobilization, consistent with the loss of latency phenotype seen in primary CD34^+^ HPC latency assays.

**Figure 3.**
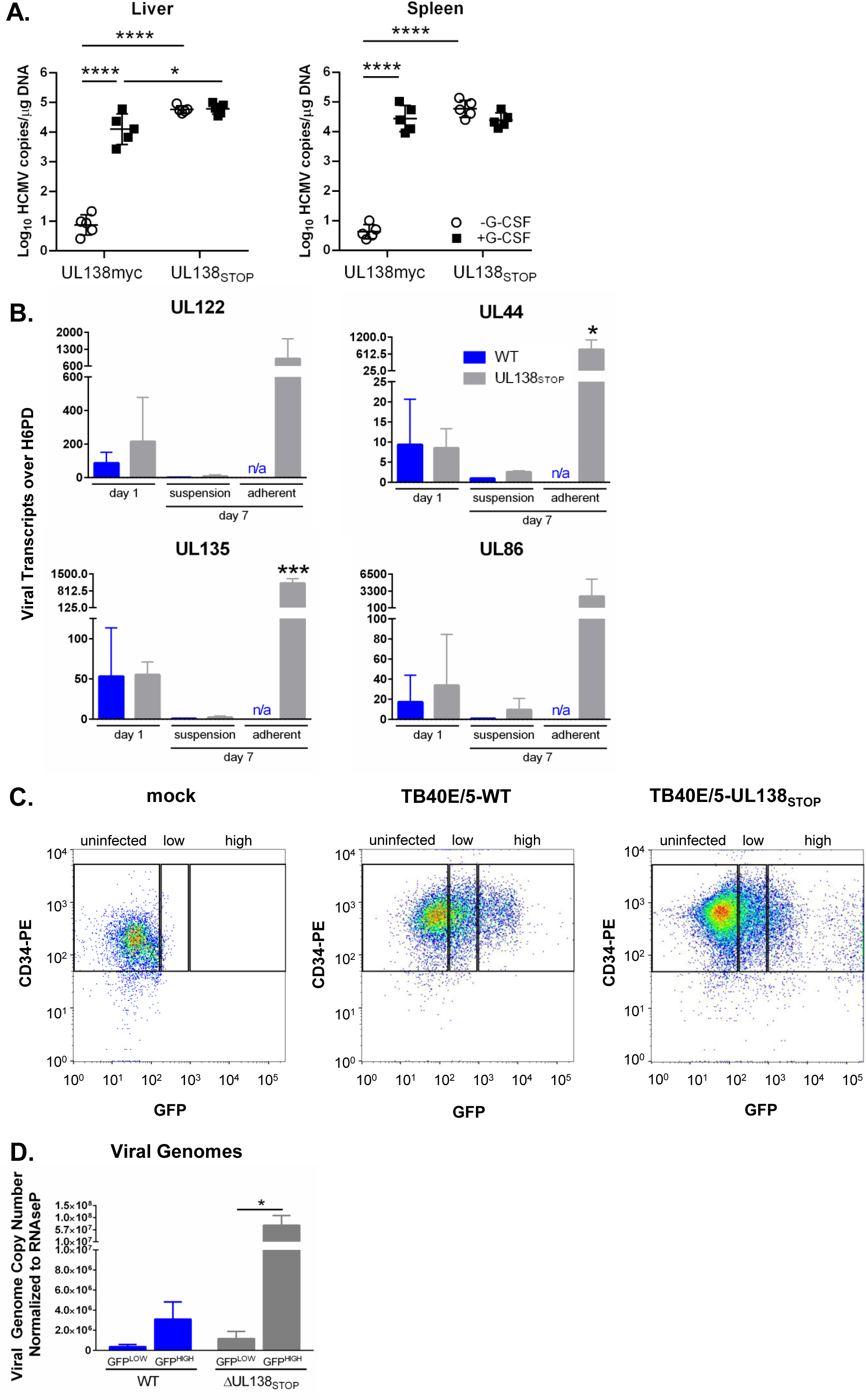
The Δ*UL138*_STOP_ loss of latency phenotype is pronounced in a subset of HCMV-infected hematopoietic cells. **A)** Humanized NSG mice (n = 10 per group) were injected with fibroblasts infected with *UL138*myc or Δ*UL138*_STOP_ HCMV. At 4 weeks post infection, half of the mice were treated with G-CSF and AMD-3100 to induce cellular mobilization and trigger viral reactivation. Control mice remained untreated. At 1 week following mobilization, mice were euthanized, and tissues were collected. Total DNA was extracted and HCMV viral load was determined by qPCR using 1 µg of total DNA prepared from liver or spleen tissue. Error bars represent standard error of the mean (SEM) between average vDNA copies from four (liver) or two (spleen) tissue sections for individual animals. All samples were compared by two-way Anova with Tukey’s multiple comparison tests within experimental groups (non-mobilized [-G-CSF] vs mobilized [+G-CSF] for each virus and between all virus groups for both non-mobilized and mobilized conditions). Statistical significance where *, P < 0.05 and ****, P < 0.00005. **B)** THP-1 cells were infected with WT or Δ*UL138*_STOP_ HCMV (MOI = 2) and cultured in suspension cell dishes for establishment of latency. Total RNA was extracted at 1 dpi from suspension cells and again at 7 dpi from suspension and adherent cells. cDNA was synthesized and viral transcripts were quantified by RT-qPCR. WT-infected cells did not spontaneously adhere to tissue culture dishes without reactivation stimulus in sufficient quantities to make cDNAs. Error bars represent SEM among three biological replicates analyzed in triplicate. Unpaired t tests were performed to compare individual time points for each virus infection by transcript. Statistical significance where *, P < 0.05 and ***, P < 0.0005. **C)** CD34^+^ HPCs were infected with WT or Δ*UL138*_STOP_ HCMV (MOI = 2) for 24 hours, then CD34/PE^+^ and GFP^+^ (infected) cells were isolated by fluorescence-activated cell sorting (FACS). WT- and Δ*UL138*_STOP_-infected populations were divided into GFP^LOW^ versus GFP^HIGH^ experimental groups, using the gating strategy shown. **D)** Pure populations of WT- or Δ*UL138*_STOP_-infected CD34^+^/GFP^LOW^ and CD34^+^/GFP^HIGH^ cells were cultured over stromal support for establishment of latency. At 10 dpi, total DNA was isolated from each experimental group and viral genomes were quantified by qPCR. Data are shown as viral genome copy number normalized to the cellular gene RNAseP. Three experimental replicates were analyzed in duplicate; error bars represent SEM among experimental replicates.

This result is inconsistent with the relatively similar patterns of viral gene expression detected between WT and Δ*UL138*_STOP_ infections in THP-1 cells (Figures 1-2). However, through the course of these studies, we observed a reproducible increase in the number of Δ*UL138*_STOP_-infected THP-1 cells that would spontaneously adhere to the cell culture dishes when compared to WT-infected THP-1s. The adherent fraction of Δ*UL138*_STOP_-infected THP-1 cells were not included in our original sequencing experiment, which was derived only from cells in suspension from each of the 216 samples. Because myeloid differentiation is linked with HCMV reactivation, we hypothesized that an increase in spontaneous differentiation in the absence of pUL138 results in increased viral gene expression, contributing to the loss of latency phenotype seen in the Δ*UL138*_STOP_ infection.

To test this hypothesis, we analyzed viral transcripts in THP-1 cells infected with WT or Δ*UL138*_STOP_ and cultured for 7 days. Total RNA was collected from suspension cells at 1 dpi during the establishment of latency. At 7 dpi, RNA was collected separately from suspension cells or adherent cells from the same dish. RT-qPCR was used to quantify viral transcripts from four genes representing 3 kinetic classes of expression (Figure 3B). Both the WT and Δ*UL138*_STOP_ infections have comparable levels of viral RNA detected at 1 dpi. By Day 7, expression of viral genes was silenced in the suspension fraction of cells infected with either WT or Δ*UL138*_STOP_ viruses. However, viral transcripts generally trend slightly higher in the Δ*UL138*_STOP_ infection, consistent with the number of viral reads detected in the sequencing data (Figures 1 and 2). In each of the three replicates, WT infection resulted in too few adherent cells in the absence of reactivation stimulus for the quantification of viral transcripts from these samples. Strikingly, in the Δ*UL138*_STOP_ infection, viral transcripts are increased 100 to 300-fold in spontaneously adherent cells relative to cells remaining in suspension and collected at the same time point. These findings indicate that the fraction of Δ*UL138*_STOP_-infected THP-1 cells that spontaneously adhere to the dish represent a distinct population that is permissive for viral gene expression in the absence of a reactivation stimulus.

We next explored this phenotype in primary CD34^+^ HPCs. GFP expressed from the SV40 early promoter in our recombinant viruses was used to detect and purify infected cells. When sorting CD34^+^/GFP^+^ cells, we observe a high GFP^+^ shift in a proportion of Δ*UL138*_STOP_-infected cells at 24 hours post infection (hpi) that is diminished in WT-infected cells (Figure 3C). We hypothesized that the GFP^HIGH^ population might represent an early readout for cells with higher levels of viral gene expression in the absence of pUL138. We collected GFP^LOW^ (middle gate) and GFP^HIGH^ (right gate) populations separately from infected CD34^+^ cells at 24 hpi, then cultured each population over stromal support as previously described (*41*) for 10 days to allow the establishment of latency. Viral genome copy number was determined by qPCR and normalized to a cellular control gene at 10 dpi (Figure 3D). When comparing the two GFP^LOW^ populations, genome copy number was approximately three-fold higher in the Δ*UL138*_STOP_-infected vs WT-infected cells. These data are consistent with an overall increase in viral replication in the absence of *UL138* during latency that is also reflected in the RNA-Seq data (Figures 1 and 2) and in the suspension fraction of THP-1 cells (Figure 3B). In the populations that were GFP^HIGH^ at 1 dpi, the number of viral genomes is almost 24 times higher in the Δ*UL138*_STOP_ infection compared to WT. This is consistent with an increase in viral transcripts in adherent THP-1 cells infected with Δ*UL138*_STOP_ (Figure 3B). Insufficient quantities of RNA were collected from the CD34^+^ HPCs to analyze viral transcripts in these cells.

Taken together, these results suggest that a distinct subpopulation of infected hematopoietic cells is present in the absence of pUL138, and that this subpopulation supports viral replication in the absence of reactivation stimuli, such as TPA or cytokine stimulation. It is possible virus replication occurs in only a fraction of Δ*UL138*_STOP_-infected cells and it is this subset that accounts for the loss of latency phenotype in primary CD34^+^ HPCs and in huNSG mice. The exclusion of the adherent fraction of THP-1 cells in our RNA-Seq analysis likely accounts for the similar viral read counts when comparing Δ*UL138*_STOP_ infection to WT infection. Further work is needed to define this subpopulation of cells and to explore a potential role for pUL138 in delaying or blocking myeloid differentiation to support the establishment and maintenance of latent infection.

### An initial burst of viral gene expression occurs in infected cells prior to silencing and is driven by the UL135 protein

Most of the variance in viral gene expression across the RNA-Seq data set is observed during infection with Δ*UL135*_STOP_ (Figure 1B), which does not reactivate from latency to produce viral progeny in CD34^+^ HPCs. This variance is most strikingly demonstrated by two aberrant patterns of viral gene expression when pUL135 is absent. The first critical difference is that the total number of viral reads is much lower during the establishment of latency (days 1 and 3) following infection with Δ*UL135*_STOP_ when compared to WT or Δ*UL138*_STOP_ in all clusters (Figure 2). The second is that viral genes belonging to cluster 4 are not efficiently expressed in the absence of pUL135 even following TPA treatment.

We hypothesized that the early detection of viral reads following infection represents an initial burst of viral gene expression that might be important for the establishment of a reactivation-competent infection. To begin to address this question, we first asked if the viral reads detected during the first 24 hours following infection were the result of *de novo* viral gene expression. THP-1 cells were pre-treated with Actinomycin D for 30 minutes to block transcription, then infected with WT or Δ*UL135*_STOP_. Total RNA was collected over the initial 24 hours of infection to assess steady-state RNA levels of viral genes from each kinetic class (Figure 4A). In infected cells pre-treated with DMSO (vehicle control), each of the representative viral transcripts increased over the first 24 hours following WT infection, whereas lower levels of viral transcripts were detected in Δ*UL135*_STOP_-infected cells (Figure 4A), consistent with the expression pattern observed in the RNA-Seq experiment (Figures 1 and 2).

**Figure 4.**
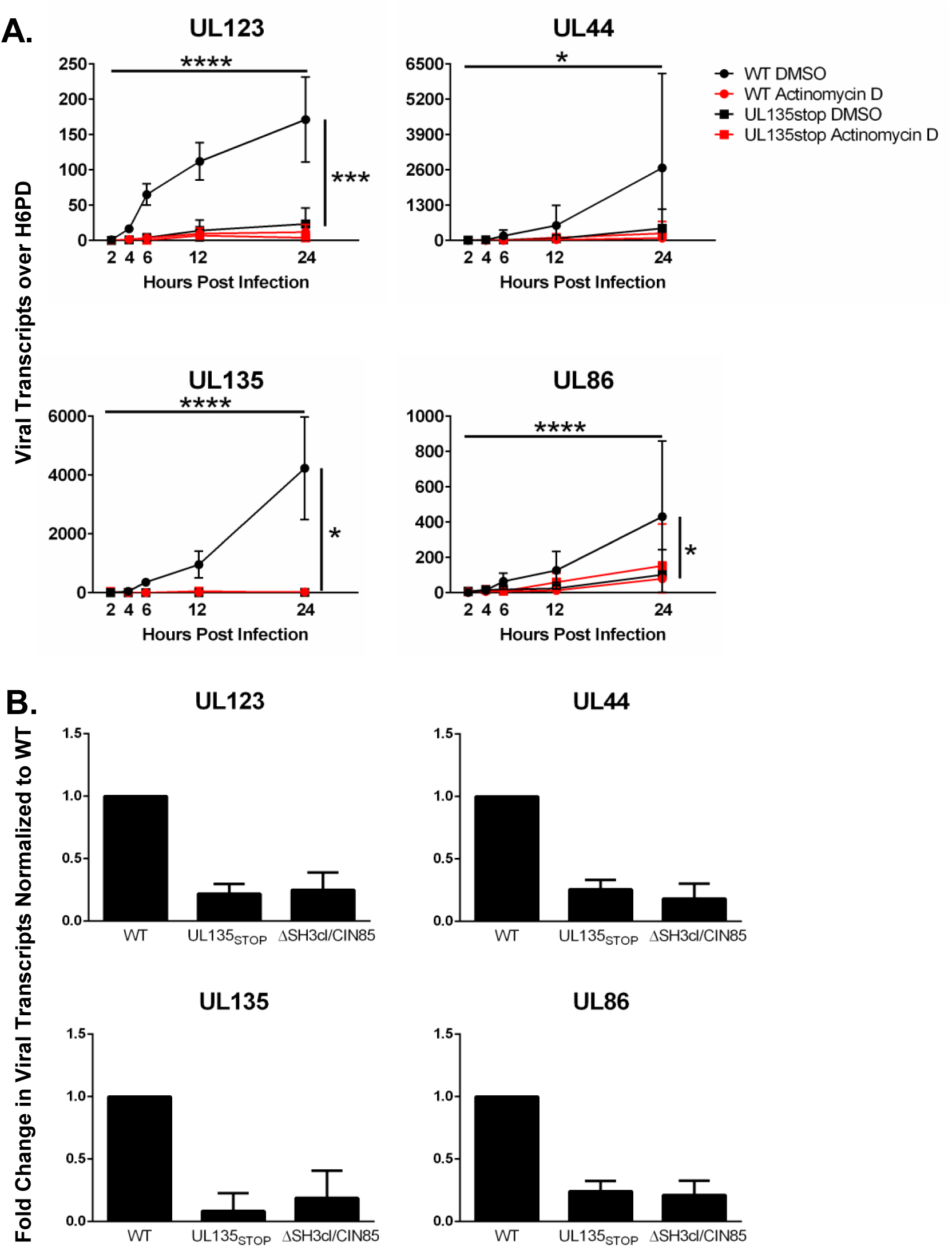
An initial burst of viral gene expression occurs in infected cells prior to silencing and is driven by the UL135 protein. THP-1 cells were pre-treated with Actinomycin D or DMSO control for 30 minutes, then infected with WT or Δ*UL135*_STOP_ HCMV (MOI = 2). Total RNA was collected over a time course of 24 hours and viral transcripts were quantified by RT-qPCR. Error bars represent SEM between three biological replicates analyzed in triplicate. All samples were compared by two-way Anova with Tukey’s multiple comparison tests across time and within experimental groups (DMSO vs Actinomycin D for each virus and WT vs Δ*UL135*_STOP_ for both DMSO and Actinomycin D treatments). Statistical significance where *, P < 0.05; ***, P < 0.0005 and ****, P < 0.00005.

When either WT-infected or Δ*UL135*_STOP_-infected cells were pre-treated with Actinomycin D, levels of viral transcripts were diminished, comparable to the levels observed in the Δ*UL135*_STOP_/DMSO treatment group (Figure 4A). Taken together, these data suggest that our detection of viral reads during the establishment of latency represents *de novo* viral gene expression and that *UL135* drives this initial burst of viral gene expression.

We previously reported that EGFR signaling promotes viral latency (*30, 42, 43*). Inhibition of EGFR or its downstream signaling pathways (PI3K/AKT or MEK/ERK) stimulates viral replication and rescues the reactivation defect in Δ*UL135*_STOP_ infection (*30, 43*). Further, a reduction in EGFR signaling due to *UL135*-mediated turnover of EGFR increases viral gene expression and reactivation from latency (*30*). *UL135* targets EGFR for rapid turnover through its interaction with cellular factors Abelson interactor 1 (Abi-1) and Cbl-interacting 85-kDa protein (CIN85) (*44*). We hypothesized that *UL135* might drive the initial burst of viral gene expression by attenuating EGFR signaling during the early hours of infection. To test this hypothesis, we infected THP-1 cells with WT, Δ*UL135*_STOP_, or a recombinant virus where the motifs required for interaction with Abi-1 and CIN85 are disrupted, ΔSH3cl/CIN85 HCMV. We collected total RNA and used RT-qPCR to compare viral transcripts from each kinetic class at 24 hpi when the initial burst of viral gene expression is first observed (Figure 4B). As expected, viral transcripts were lower in the Δ*UL135*_STOP_ infection when compared to WT infection.

Transcripts detected in ΔSH3cl/CIN85 infection were comparable to the levels observed in Δ*UL135*_STOP_ infection. These results suggest that pUL135 interaction with Abi-1 and CIN85 is required for the initial burst of viral gene expression, indicating a potential role for attenuation of EGFR signaling. Ongoing studies are aimed at examining the specific mechanisms driving the initial burst of viral gene expression and whether it is required for HCMV reactivation from latency.

### Motif analysis reveals candidate transcription factors for controlling expression of UL*b*’ genes

The strikingly low read counts observed for cluster 4 viral genes in Δ*UL135*_STOP_ infection (Figure 2) is perhaps one of the most surprising results from our analysis. The eleven genes in cluster 4 reside in the UL*b*’ region of the HCMV genome. UL*b*’ is present in clinical isolates and low-passage strains of HCMV but is consistently lost in laboratory-adapted strains following successive passaging through replication permissive cell lines. The UL*b*’ gene region spans *UL133* through *UL150* and encodes proteins with roles in immune evasion, viral dissemination in the host, and/or modulating latency and reactivation. Notably, this ∼15 kb of the genome includes the *UL133-UL138* locus encoding *UL135* and *UL138*. Because expression of a block of UL*b*’ genes is diminished in the absence of pUL135, our data suggested that *UL135* functions as a master regulator of this locus, controlling expression levels of at least eleven viral genes.

To confirm the results of our transcriptome analysis, we performed RT-qPCR to quantify expression levels of two representative UL*b*’ genes during WT, Δ*UL135*_STOP_, and Δ*UL138*_STOP_ infection of THP-1 cells (Figure S2A). In the WT and Δ*UL138*_STOP_ infections, both *UL135* and *UL138* transcripts were expressed at early time points, then decreased during the latency period. The transcripts were induced again following TPA treatment. As expected, expression of the viral genes is increased in the Δ*UL138*_STOP_ infection relative to the WT infection. Consistent with the transcriptome data (Figure 2), *UL135* and *UL138* transcripts are expressed at very low levels across the time course in Δ*UL135*_STOP_ infection (Figure S2A).

To ensure that the Δ*UL135*_STOP_ virus was competent to express these UL*b*’ genes, we analyzed their expression following infection in MRC-5 fibroblasts, a model for productive virus replication. In contrast to THP-1 cells, both *UL135* and *UL138* transcripts are expressed to near wildtype levels in Δ*UL135*_STOP_ infection of fibroblasts (Figure S3B). Taken together, these data suggest that the differences in transcription in the absence of *UL135* are due to a cell type-specific role in viral gene transcription rather than a loss of the UL*b*’ gene locus in these experiments.

The functions thus far defined for pUL135 are achieved via modulation of cellular signaling pathways (*30, 44, 45*). We therefore hypothesized that *UL135* regulates transcription of the UL*b*’ locus via an indirect mechanism such as the regulation of cellular transcription factors. Accordingly, we used a bioinformatics approach to identify cellular transcription factors that are predicted to regulate gene expression in the UL*b*’ locus and whose expression is altered depending on the presence of pUL135. We used the simple enrichment analysis (SEA) algorithm to identify transcription factor binding sites that are enriched in each of our four viral gene expression clusters. In cluster 4 (UL*b*’ genes dependent on pUL135 for their expression), our analysis uncovered significant enrichment of binding sites for twenty-five cellular transcription factors (Supplementary Data Set 1). We then used differential expression analysis to identify which of these transcription factors are regulated at the transcript level in response to pUL135 (Supplementary Data Set 2) by comparing infections with pUL135 present (WT and Δ*UL138*_STOP_ averaged) or absent (Δ*UL135*_STOP_). These analyses resulted in the identification of nine transcription factors that are predicted to regulate cluster 4 viral genes and are differentially expressed when pUL135 is absent in infection (Figure 5A).

**Figure 5.**
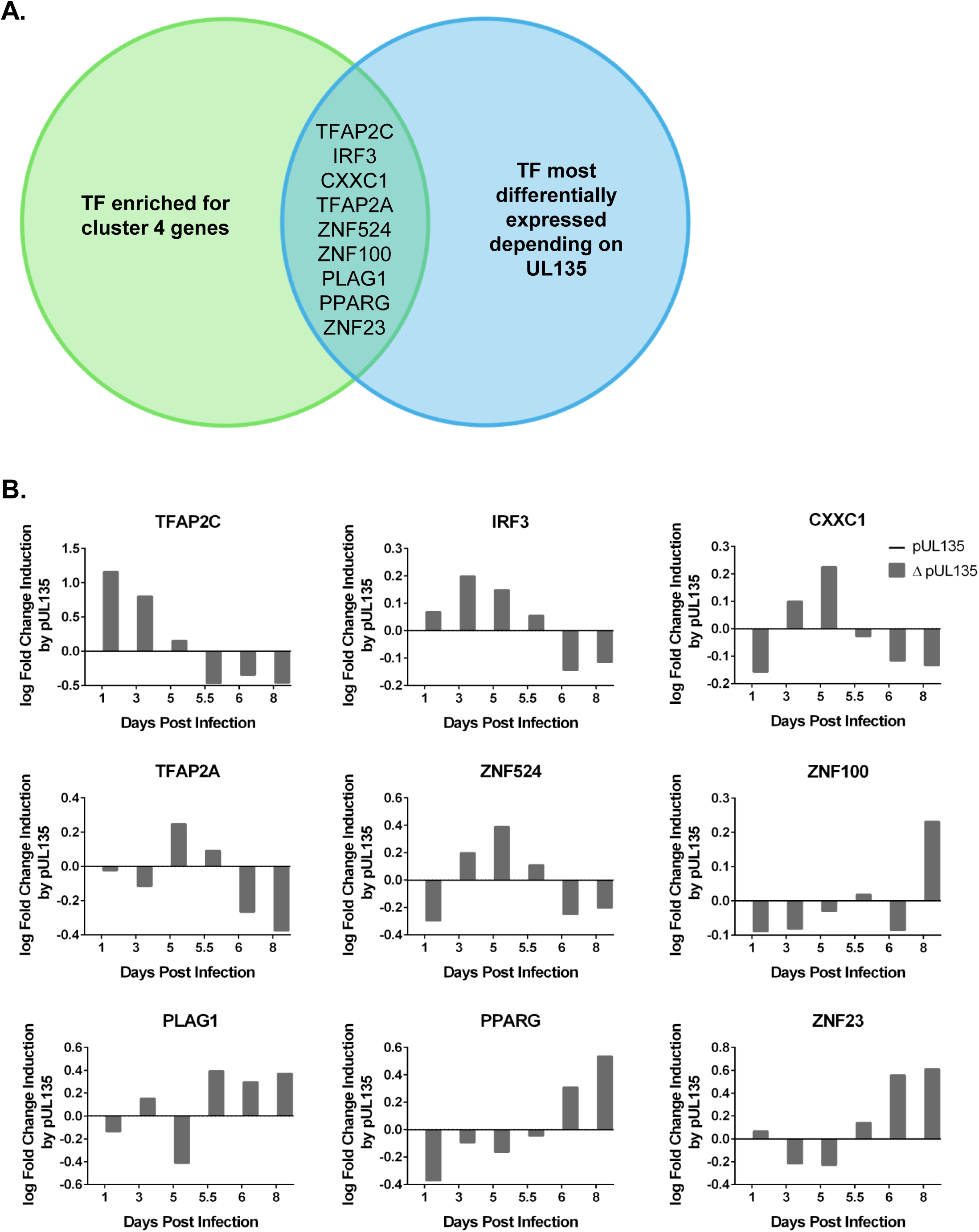
Motif analysis reveals candidate transcription factors driving expression of UL*b*’ genes. **A)** Graphical representation of motif analysis. A simple enrichment analysis (SEA) (*66*) was performed to identify predicted transcription factor binding motifs that are enriched in cluster 4 genes compared to the total HCMV genome (Supplementary Data Set 1). These transcription factors were then ranked by degree of differential expression at each time point dependent on the presence of pUL135 in our RNA-Seq analysis (Supplementary Data Set 2). When compared, these analyses generated a list of nine transcription factors that are regulated by pUL135 and are significantly more likely to control gene expression from the cluster 4 genes. **B)** RNA expression profiles of each of the nine candidate transcription factors from the RNA-Seq data set are shown. Numerical values are log2 fold change as a function of read count and represent the average of four biological replicates sequenced per experimental group. Data are normalized to show log2 fold change in expression when pUL135 is present (grey bars; average of WT and Δ*UL138*_STOP_ infection at each time point) over absence of pUL135 (Δ*UL135*_STOP_ infection). Values for Δ*UL135*_STOP_ infection are set to zero so that an induction or repression of transcripts corresponds to a positive or negative number, respectively.

We next used our RNAseq data to plot the log fold change in expression of each candidate transcription factor when pUL135 is present (Figure 5B). The bars represent the log2 fold change in expression of each transcription factor in *UL135*-expressing infection relative to Δ*UL135*_STOP_ infection (Figure 5B). Data are shown for Days 1, 3 and 5 post infection as latency is established, and for Days 5.5, 6, and 8 following reactivation stimulus (Figure 5B). Of the nine transcription factors identified by our motif analysis, three are induced by *UL135* following TPA treatment: Pleomorphic adenoma gene one (PLAG1), Peroxisome proliferator-activated receptor gamma (PPARγ), and Zinc finger protein twenty-three (ZNF23). Future work will examine the potential contribution of each of these transcription factors in driving re-expression of the ULb’ viral genes in concert with pUL135 and whether re-expression of these genes is key for reactivation from latency.

## Discussion

The development of next generation sequencing techniques has provided a valuable framework for understanding challenging and complex transcriptomes during viral infection. RNA-Seq technology allows for comprehensive analysis of both host and pathogen gene expression across the course of infection. Here, we have harnessed this technology to assemble a complete representation of viral gene expression in the THP-1 model of HCMV latency and reactivation. By comparing viral gene expression among the wildtype virus and two recombinant viruses missing either the gene required to establish latency (*UL138*), or the gene required for reactivation from latency (*UL135*), our data reveal novel intricacies of the viral gene expression program controlling the switch between latent and replicative infection.

This study has revealed clear transcriptional switches associated with the establishment, maintenance, and exit from latency. Infection of THP-1 cells results in an initial burst of viral gene expression that is broad and is uncoupled from the orderly progression of viral gene expression (immediate early, early, late phases) seen during productive infection. A similar phenomenon has been described for Herpes simplex virus 1 (HSV-1) during the earliest stages of reactivation from latency (*46–48*). Broad and disordered HSV-1 gene expression occurs during the “animation” phase and is proposed to give rise to the viral transactivator VP16 so that replication can proceed in a coordinated kinetic cascade during the “synthesis” phase. In our data set, most viral genes are silenced to very low levels of expression following the initial burst, representative of the establishment and maintenance of latency. Following a TPA reactivation stimulus, viral genes are re-expressed to increased levels relative to those immediately following infection.

CD34^+^ primary HPCs, the gold standard for HCMV latency, are a heterogeneous population containing cells at various stages of differentiation and lineage commitment, which complicates transcriptome studies. Although the CD34^+^ HPC model likely reflects the true nature of a more dynamic persistence *in vivo*, we are limited not only by availability, but also by the ability to achieve a coordinated entry into and synchronous and robust exit from latency.

Additionally, the CD34^+^ HPCs used in our previous transcriptome study produce very low levels of viral transcripts and the clinical samples used in the same study typically have viral carriage between 1 in 10,000 to 1 in 25,000 cells (*11*). Because triggering a synchronous reactivation is not possible in the CD34^+^ HPC model, this work was also limited to latency time points only.

We used the THP-1 cell line to compensate for the challenges presented by CD34^+^ HPCs and clinical samples such as their heterogeneity and donor variability, exceptionally low level of viral transcripts, and lack of a good solution for including reactivation time points. In addition to expanding our series of time points to include a synchronous reactivation from latency, use of the THP-1 cell line allowed us to perform this study using four biological replicates to optimize statistical power. However, cell line models are limited in their ability to faithfully recapitulate every aspect of latency and reactivation. For example, THP-1 cells cannot efficiently synthesize HCMV genomes nor replicate productively, even following differentiation with TPA and re-expression of viral genes (*22, 31, 34*). As such, THP-1 cells are an effective tool for examining viral gene expression patterns in response to reactivation stimuli but cannot be used to assess a true reactivation from latency as defined by the production of viral progeny. Despite their inherent differences, the two models together provide a more complete picture of latency and reactivation. Indeed, many of the results from our CD34^+^ HPC model were recapitulated in the THP-1 model. Both studies are consistent with latency transcriptomes produced by other groups (*12–14*) in showing broad viral transcription during latency, but with lower levels of transcripts than in lytic infection. Additionally, *UL5*, *UL40*, *UL22A*, *RL12*, *RL13*, *UL4*, *UL78*, *UL44*, *UL132*, *US18*, and *UL148* were among the highest expressed viral genes in both the current study and in the primary CD34^+^ HPC study (*11*). Because these genes are expressed to high levels during both latency and reactivation, they may not play a role in regulating the transition between the two infectious states. In addition to defining contextual changes in viral gene expression patterns, the current study revealed important roles for both pUL138 and pUL135 in regulating viral gene expression and potentially cellular differentiation to navigate the transition between latent and replicative states of infection.

*UL138* has a well-defined role promoting the establishment and maintenance of latency in both the CD34^+^ HPC model (*27, 29, 30, 40*) and the humanized mouse model (Figure 3A). It was therefore unexpected that viral gene expression would be silenced in Δ*UL138*_STOP_ infection relative to the wildtype infection (Figures 1 and 2). However, further investigation revealed a distinct population of hematopoietic cells within the infected population that support a more replicative infection in the absence of *UL138* and seem to account for the Δ*UL138*_STOP_ loss-of-latency phenotype. This manifests in THP-1 cells that adhere to the tissue culture dish and express very high levels of viral transcripts in the absence of a reactivation stimulus (Figures 3B and 3C) which were not captured in the transcriptome. Analogous findings are also seen in CD34^+^ HPCs infected with Δ*UL138*_STOP_ in that a subset of infected cells express inordinately increased levels of GFP (a proxy for viral gene expression) as early as 24 hpi and have more viral genomes per cell at 10 dpi relative to WT infection, suggesting a more replicative infection where latency is not established (Figures 3D and 3E). Because this phenotype appears only in Δ*UL138*_STOP_ infection and only in a small subset of infected cells, it appears that both viral expression of *UL138* and some aspect of the cellular environment that varies in both the THP-1 and CD34^+^ HPC models are important for the establishment of latency.

A recent single cell sequencing study (*14*) identified intrinsic expression of interferon-stimulated genes (ISGs) as the strongest predictor for a replicative versus non-replicative infection outcome. This study found that intrinsic ISG expression correlates with the differentiation state of cells, where monocytes express the highest levels of ISGs, followed by macrophages, and then fibroblasts with the lowest ISG expression. At the same time, monocytes are the least permissive for replicative HCMV infection, followed by macrophages, and then fibroblasts. The authors concluded that high levels of ISGs lead to a non-productive infection while low levels of ISGs support a productive infection. Importantly, they showed that intrinsic, but not induced, levels of ISGs were critical for curbing viral gene expression and these levels were determined by IRF9 and STAT2 (*14*). It follows that as HCMV pushes differentiation of an infected cell along the myeloid lineage, it would have lower basal levels of ISG expression and allow for more viral gene expression. Therefore, an additional mechanism must exist to keep ISG levels high and viral gene levels low for the establishment and maintenance of latency. We have shown that *UL138* interacts with a UAF1-USP1 complex to sustain STAT1 activation and enhance an early ISG response that restricts viral replication (*40*). Taken together, these studies suggest that both conditions (high intrinsic levels of ISGs and expression of pUL138 during infection) must be satisfied for efficient establishment of latency. In this scenario, the non-adherent THP-1 cells and GFP^LOW^ CD34^+^ cells would intrinsically express high levels of ISGs, leading to decreased viral gene expression for the establishment of latency. In the WT infection, *UL138* would enhance and sustain an ISG response to facilitate this process, leading to fewer cells that are productively infected. In contrast, the fraction of THP-1s that adhere spontaneously and the CD34^+^ cells with higher levels of GFP following infection with the Δ*UL138*_STOP_ virus might endure a “double-hit” of having lower intrinsic levels of ISG expression that cannot be overcome in the absence of pUL138. These differences cannot be evaluated in our existing transcriptome data which only includes the *UL138*_STOP_-infected suspension fraction of THP-1 cells. Additional work is needed to fill important gaps in our knowledge. It will be critical to identify the cellular factors that differ in hematopoietic cells that are predisposed to a more replicative infection versus those that support a latent infection, the specific mechanisms that drive those differences, as well as the potential role of pUL138 in stalling myeloid differentiation of infected cells to promote a latent infection.

The current study also expanded our understanding of *UL135* as a driver of replicative HCMV infection. Our results demonstrate that pUL135 i) drives an initial burst of broad viral gene expression in the early hours of infection and ii) functions as a master regulator of viral gene expression from the UL*b*’ gene region encoding the UL133-UL138 proteins that function to modulate latent versus replicative infection in hematopoietic cells. These findings suggest two temporally (and perhaps mechanistically) distinct strategies for driving broad viral gene expression prior to the establishment of latency versus re-expression of select viral genes following a reactivation stimulus. We have shown that the initial burst of viral gene expression requires the interaction between *UL135* and Abi-1 and CIN85, which we have shown directs EGFR for turnover in infection, as either infection with recombinant viruses lacking *UL135* or expressing a variant of *UL135* where motifs required for interaction with Abi-1 and CIN85 have been disrupted results in a diminished initial burst. It is possible that alterations in EGFR signaling over the early course of infection change the balance of transcription factors that would drive viral gene expression during the initial burst. For example, previous work in our lab has identified EGR1 as a transcription factor that is up-regulated via EGFR signaling, then binds the HCMV genome to drive expression of the latency determinant *UL138* to promote silencing during latency (*43*). Future work will identify transcription factors that are responsive to EGFR signaling and assess their potential to drive broad viral gene expression during the initial burst. Importantly, the initial burst is absent only during infections where *UL135*, which is required for reactivation from latency, is not expressed (Figures 1 and 2) or is prevented from turning over EGFR from the cell surface (Figure 4B). These data suggest an important link between the initial burst of viral gene expression, EGFR signaling, and subsequent viral reactivation from latency. Given the ability of early events during alphaherpesvirus infection to affect the re-expression of viral genes during reactivation (*49*), we hypothesized that the initial burst of viral gene expression is required to optimize infection conditions to support a successful reactivation from latency. Future work will test whether the initial burst is important for a robust reactivation from latency and focus on identifying the cellular conditions required to support viral reactivation.

In contrast to the broad pattern of viral gene expression contributing to the initial burst, the role of pUL135 in the re-expression of viral genes following a reactivation stimulus is more focused. The pUL135-dependent response is limited to eleven viral genes from the UL*b*’ genomic region, and all other viral genes are re-expressed in the absence of pUL135. Although high expression levels of the pUL135-independent genes could be the result of an overwhelming response to TPA treatment, it is nonetheless clear that the UL*b*’ genes are regulated differently than the more TPA-responsive genes and that pUL135 is required for their expression. The *UL133*-*UL138* latency locus encodes at least four proteins that modulate viral replication to promote latency or reactivation, and these are among the eleven genes that are dependent on pUL135 for their re-expression following reactivation stimulus. The discovery of pUL135 as a driver of gene expression from the *UL133*-*UL138* locus is consistent with our previous work showing that the 33 kDa isoform of pUL136 is important for driving reactivation from latency (*50, 51*) and that stabilization of this isoform overcomes the requirement for pUL135 in promoting viral replication in hematopoietic cells (*52*).

Simple enrichment analysis and differential expression analysis identified transcription factors that were differentially expressed at the RNA level depending on presence of pUL135 and that also have an enrichment of predicted DNA binding sites in the UL*b*’ genomic region. Two of these, PLAG1 and PPARγ, are associated with growth factor signaling pathways and could link the initial burst of viral gene expression with re-expression of crucial UL*b*’ genes through a similar, although more targeted, mechanism for pUL135 regulation of viral gene expression. The PLAG1 transcription factor targets numerous genes encoding growth factors and growth factor receptors (*53*) which could alter these cell signaling pathways to support full re-expression of HCMV genes. Additionally, chemical inhibition of EGFR signaling results in induction and nuclear accumulation of PPARγ (*54, 55*) which could then drive transcription of UL*b*’ genes in addition to its cellular targets. Future work will determine the contribution of each candidate transcription factor in driving gene expression of the eleven cluster 4 genes and then dissect the molecular mechanisms involved including expression kinetics, localization, and activation of the transcription factors.

The polycistronic *UL133*-*UL138* locus encodes determinants of HCMV latency and reactivation (*26*). The relative accumulation of pUL135 and pUL138 in infected hematopoietic cells is likely critical in dictating the outcome of infection as replicative or relatively silenced. Here, we have revealed roles for *UL138* and *UL135* in the regulation of HCMV gene expression and potentially hematopoietic cell differentiation to navigate the switch between latent and replicative infection. Unraveling the mechanistic basis for these functions and identifying crucial cellular interactors will deepen our understanding of the molecular events regulating HCMV latency and reactivation and provide potential therapeutic targets for controlling HCMV infection.

## Materials & Methods

### Data Availability

The data set that supports this study has been deposited into the Gene Expression Omnibus (GEO) database under the following accession code GSE266854.

### THP-1 monocyte model for latency and reactivation

THP-1 cells were purchased from ATCC (Manassas, VA) and cultured in Roswell Park Memorial Institute (RPMI) 1640 medium (Cytiva Hyclone, Marlborough, MA) supplemented with 10% fetal bovine serum (FBS) (Gibco Thermo Fisher, Waltham, MA), 2 mM L-alanyl-glutamine (Corning, Corning, NY), 0.05 mM β-mercaptoethanol (Sigma-Aldrich, St. Louis, MO), and 100U/mL penicillin -100 µg/mL streptomycin (Gibco Thermo Fisher). THP-1 cells were infected as monocytic suspension cells at a density of 5 x 10^5^ cells per mL in RPMI cell culture media. Stocks of TB40/E HCMV expressing green fluorescent protein (GFP) were titrated using THP-1 cells so that infections were carried out to result in 40-60% GFP-positive cells at 24 hours post infection. A multiplicity of infection (MOI) of 2 plaque forming units per cell, as determined by TCID_50_ in MRC-5 fibroblasts, was used as a starting titration. Infected cell suspension was mixed by periodic rocking in untreated six well plates designed for suspension cells (Sarstedt, Nümbrecht, Germany), then a spinoculation was performed by centrifugation at 450 x g for 20 minutes. Cells were cultured for 5 days post infection (dpi) and concentration was maintained between 4 x 10^5^ and 8 x 10^5^ cells/mL by adding cell culture media. On day 5, cells from each experimental group were pooled and pelleted at 120 x g for 7 minutes, then resuspended at 5 x 10^5^ cells/mL. Cells were treated with 100 nM 12-O-Tetradecanoylphorbol-13-acetate (TPA) (LC Laboratories, Woburn, MA) and plated on tissue culture-treated plates to trigger monocyte-to-macrophage differentiation and viral reactivation or treated with an equivalent volume of dimethyl sulfoxide (DMSO) (Sigma-Alrich) solvent control and cultured in untreated six well plates as described above. Total RNA was collected at the indicated time points during infection, as described below in *Reverse Transcriptase quantitative polymerase chain reaction (RT-qPCR)*.

### MRC-5 fibroblast model for replicative infection

MRC-5 human embryonic lung fibroblasts were purchased from ATCC (Manassas, VA) and cultured in Dulbecco’s Modified Eagle Medium (DMEM) (Gibco Thermo Fisher) supplemented with 10% FBS (Gibco Thermo Fisher), 10 mM HEPES (Corning), 2 mM L-alanyl-glutamine (Corning), 1 mM sodium pyruvate (Gibco Thermo Fisher), 0.1 mM non-essential amino acids (Gibco Thermo Fisher), and 100 U/mL penicillin -100 µg/mL streptomycin (Gibco). MRC-5 cells were infected with TB40/E-5 HCMV (MOI = 1). At 2 hours post infection (hpi), virus inoculum was removed and replaced with fresh

DMEM cell culture media. Total RNA was collected at the indicated time points during infection, as described below in *Reverse Transcriptase quantitative polymerase chain reaction (RT-qPCR)*.

### Viruses

The TB40/E-5 bacterial artificial chromosome (BAC) was previously engineered to express green fluorescent protein (GFP) as a soluble marker for infection (*28, 56*). The Δ*UL135*_STOP_ and Δ*UL138*_STOP_ recombinants were made from the parental wildtype (WT) BAC as previously described (*29*). For each of these recombinant viruses, ATG start codons were mutated to TAG stop codons. The Δ*UL135*_STOP_ recombinant was mutated at amino acid positions M1, M21, and M97 to abrogate expression of the pUL135 protein, and therefore cannot reactivate from latency. The Δ*UL138*_STOP_ recombinant was mutated at amino acid position M1 and does not express the pUL138 protein, rendering it defective for establishment of latency. The *UL138*myc recombinant was made from the parental WT BAC as previously described (*57*) by cloning the myc epitope tag in frame onto the C-terminus of *UL138*. The ΔSH3cl/CIN85 recombinant (*44*) was previously made by incorporating alanine substitutions of key amino acid residues that mediate the interaction of pUL135 with proteins containing a Src homology 3 (SH3) domain. These mutations produced a virus where pUL135 is expressed but cannot interact with Abelson interactor 1 (Abi-1) and Cbl-interacting 85-kDa protein (CIN85) to regulate epidermal growth factor receptor (EFGR).

### RNA isolation, NGS library preparation, and sequencing

RNA was extracted from THP-1 cells at Days 1, 3, 5, 5.5, 6, and 8 following infection (or mock-infection) for transcriptomic profiling. RNA was isolated with a Quick-DNA/RNA™ Miniprep kit (Zymo, Irvine, CA) then treated with 5U/sample DNAseI (Zymo) and processed with an RNA Clean & Concentrate kit (Zymo). The next-generation sequencing (NGS) library was constructed by the University of Arizona Genomics Core (UAGC) facility. RNA integrity was assessed by capillary gel electrophoresis using a fragment analyzer (Agilent, formerly Advanced Analytical Technologies, Santa Clara, CA) and measured through RNA Integrity Number (RIN) score (mean score = 9.0). Presence of residual genomic DNA (gDNA) was also assessed by this method. cDNA libraries were prepared with an Illumina TruSeq Stranded mRNA kit (Illumina, Inc, SanDiego, CA) and a KAPA Dual-Indexed Adaper kit (KAPA Biosystems, Wilmington, MA). The NGS library was quantified with a qPCR-based KAPA Library Quantification kit (Roche, Basel, Switzerland).

Samples were sequenced at the University of California San Francisco (UCSF) Center for Advanced Technology, using the NovaSeq 6000 platform (Illumina, Inc). Paired-end sequencing with a 150 bp read length was performed on 216 samples loaded onto an S4 flow cell, with 72 samples per lane. Base calling was performed with the Real Time Analysis (RTA3) software from Illumina.

### RNA-Seq data preprocessing and analysis

Raw reads quality was assessed using FastQC 0.1 (*58*) and reads were trimmed using Trimmomatic 0.39 (*59*). Principal Component Analysis (PCA) was carried out with the ggplot2 package (*60*). Reads were aligned to combined (concatenated) human reference genome GRCh38 (ensemble version 98) and human herpesvirus 5 strain TB40/E clone TB40-BAC4 using STAR aligner (*61*). Alignment ratio was similar for all samples and the mean percentage of uniquely mapped read counts was 93.23%. Gene-level counts were determined using featureCounts function from Rsubread (*62*). Genes with more than 0.6 CPM (counts per million) in at least two samples were retained for further analysis. Gene-level count data were normalized using the *voom* method from limma (*63*). Normalized expression data of viral genes was utilized for clustering via k-means approach (k = 4). Visual assessment of the expression signature of each recombinant virus group was performed via heatmap with the *ComplexHeatmap* package from R (*64, 65*). Viral gene cluster enriched transcription factor motifs were detected by using the *SEA* (Simple Enrichment Analysis) method (*66*) against the CIS-BP database of transcription factors and their DNA binding motifs from *MEME Suite* (*67*). Differential expression analysis between different groups of recombinant viruses was carried out through negative binomial modeling of gene expression with the *DESeq2* package from R (*68*).

### Reverse Transcriptase quantitative polymerase chain reaction (RT-qPCR)

Total RNA was extracted using a Quick-DNA/RNA™ Miniprep kit (Zymo), then treated with 5U/sample DNAseI (Zymo) and processed with an RNA Clean & Concentrate kit (Zymo) according to the manufacturer’s protocol. cDNA was synthesized using the Transcriptor First Strand cDNA Synthesis Kit (Roche). Briefly, total RNA (400ng) was combined with 2.5 µM anchored-oligo(dT)18 Primers and denatured at 65°C for 10 minutes. A Reverse Transcriptase (RT) master mix (1x Transcriptor RT Reaction Buffer, 40 U/µL Protector RNase Inhibitor, 10 mM Deoxynucleotide Mix, 20 U/µL Transcriptor RT) was added to the template-primer mix. A no Reverse Transcriptase (RT-) control was made by substituting water for RT in a single reaction. Samples were incubated in a Mastercycler® X50 (Eppendorf, Hamburg, Germany) for 60 minutes at 50°C, then raised to 85°C for 5 minutes to inactivate the Reverse Transcriptase. Final reaction products were diluted 1:4 in PCR grade water to reduce salt concentrations, and the resulting single-stranded cDNA was amplified by quantitative polymerase chain reaction (qPCR). The LightCycler® 480 (Roche) was used to amplify cDNAs in a mix of 1x Lightcycler® 480 SYBR Green I Master Mix (Roche) and 0.2 µM of a series of sequence-specific primer pairs (see Table 1 for detailed target sequences). Relative expression of each mRNA was calculated using the Pfaffl method, which allows for adjustments based on the efficiency of individual primer pairs (*69*) and increases accuracy when comparing relative expression of multiple genes. Primer efficiencies were calculated using an internal standard curve of cDNA made from lytically infected fibroblasts collected at multiple time points and pooled to include viral gene expression from each kinetic class.

**Table 1.**
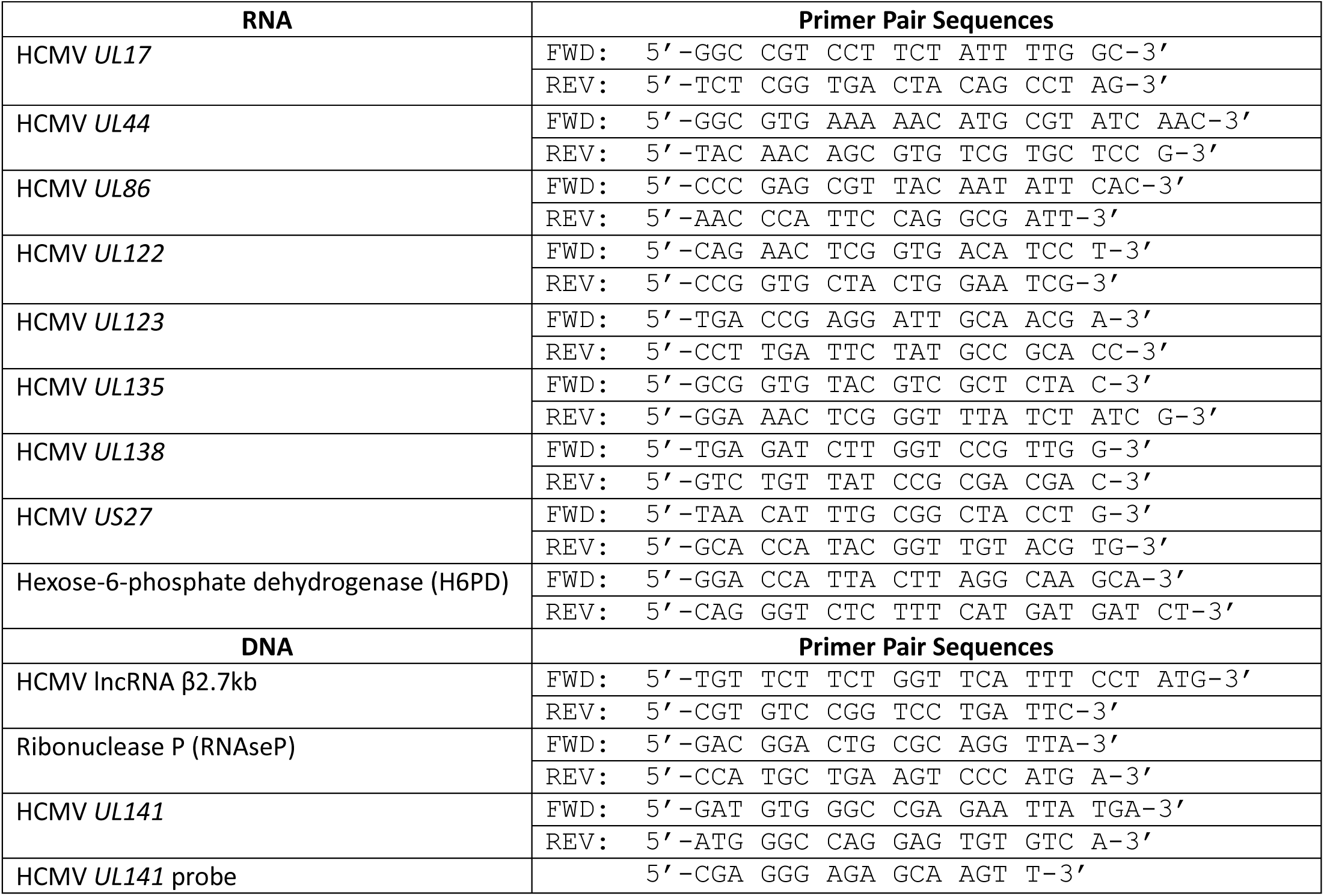
Primer sequences used in this study. Sequences are shown for all primers used in this study. Primer pairs labeled RNA were used for RT-qPCR to quantify viral transcripts. Primers labeled DNA were used for qPCR to quantify viral genome copy numbers.

### Engraftment and Infection of huNSG mice

All animal studies were carried out in strict accordance with the recommendations of the American Association for Accreditation of Laboratory Animal Care (AAALAC). The protocol was approved by the Institutional Animal Care and Use Committee (protocol 0922) at the Vaccine and Gene Therapy Institute at Oregon Health and Sciences University (OHSU). NOD-*scid* IL2Rγ_c_^null^ mice of both sexes were maintained in a pathogen-free facility at OHSU. Humanized mice were generated as previously described (*70*). At 12-14 weeks post engraftment, the animals were treated with 1 mL of 4% thioglycolate (Brewer’s medium; BD) via intraperitoneal (IP) injection to recruit monocytes/macrophages. After 24 hours, mice were inoculated with TB40/E-*UL138*_myc_ or TB40E-Δ*UL138*_STOP_-infected fibroblasts (approximately 10^5^ PFU per mouse) via intraperitoneal (IP) injection. A control group of engrafted mice was mock-infected using uninfected fibroblasts. The virus was reactivated as previously described (*70*). Briefly, half of the mice were treated with G-CSF and AMD-3100 at 4 weeks post infection to induce cellular mobilization and trigger viral reactivation. Control mice remained untreated. At 1 week post mobilization, mice were euthanized, and tissues were collected. Total DNA was extracted and HCMV viral load was determined by qPCR using 1 µg of total DNA prepared from liver or spleen tissue.

### Cell Sorting for CD34^+^ HPC Latency Culture

CD34^+^ human progenitor cells (HPCs) were obtained from bone marrow harvests; either from de-identified medical waste at Banner - University Medical Center on the University of Arizona campus or purchased from AllCells (Alameda, CA). CD34^+^ HPCs were isolated using a CD34 MicroBead kit according to manufacturer’s instructions (magnetically activated cell sorting or MACS; Miltenyi Biotec, San Diego, CA). Pure populations of CD34^+^ HPCs were infected with TB40/E-WT or TB40/E-Δ*UL138*_STOP_ HCMV (MOI = 2) expressing GFP as a marker for infection. At 24 hpi, infected (GFP^+^) CD34^+^ cells were isolated by fluorescence activated cell sorting (FACS) and collected as separate GFP^LOW^ and GFP^HIGH^ populations. CD34^+^ HPCs were then cultured in Myelocult H5100 (Stem Cell Technologies, Cambridge, MA) supplemented with hydrocortisone, 100 U/mL penicillin, and 100 µg/mL streptomycin and maintained in long-term co-culture with M2-10B4 and S.l./S.l. murine stromal cell lines (Stem Cell Technologies, Vancouver, Canada) to establish and maintain viral latency (*41*). Total DNA was collected at 10 dpi and viral genome copy number in each subset of latently infected cells was determined by qPCR.

### Quantitative polymerase chain reaction (qPCR) for measuring viral genomes

For the latently infected CD34^+^ HPCs, total DNA was isolated using a Quick-DNA/RNA™ Miniprep kit (Zymo) according to the manufacturer’s protocol. Absolute viral genome copy number was calculated by quantitative polymerase chain reaction (qPCR) using primers targeted against the genomic region corresponding to the non-coding HCMV β2.7 RNA. The number of viral genomes present in each sample was quantified relative to BAC standard curve. Each sample was then normalized to the cellular gene Ribonuclease P (RNAseP). For the huNSG mice, total DNA was isolated from liver and spleen tissue using the DNAzol method (Life Technologies, Carlsbad, CA) according to the manufacturer’s directions. Primers and a probe recognizing HCMV *UL141* were used to quantify HCMV genomes. Viral genomes in humanized mice were normalized to 1mg input DNA. Primer sequences are shown in Table 1.

### Inhibitors

Actinomycin D is a general inhibitor of transcription that intercalates into DNA and blocks RNApol II activity. THP-1 cells were pre-treated with 0.1 µg/mL of Actinomycin D (Sigma-Aldrich, St, Louis, MO) for 30 minutes, then infected with TB40/E-WT or TB40/E-Δ*UL135*_STOP_ HCMV (MOI = 2). Total RNA was collected over a time course of 24 hours and isolated using a Quick-DNA/RNA™ Miniprep kit (Zymo) according to the manufacturer’s protocol. Viral transcripts from each kinetic class were quantified by RT-qPCR to monitor *de novo* viral gene expression during the initial burst. Primer sequences are shown in Table 1.

## Acknowledgments

The authors wish to acknowledge Jonathan Galina-Mehlman and the Arizona Genetics Core (AZGC) for NGS library preparation and technical advice. We are grateful to Dr. Emmanuel Katsansis and the Banner-University Medical Center stem cell team for medical discards of bone marrow for primary CD34^+^ HPCs. We thank Mark Curry and the University of Arizona Cancer Center Flow Cytometry and Human Immune Monitoring Shared Resource for assistance with fluorescence-activated cell sorting (FACS) of infected CD34^+^ HPCs for long-term culture. We thank Matt Huntley for assistance in collecting and analyzing data in MRC-5 fibroblasts.

---

**Supplementary Figure 1.**
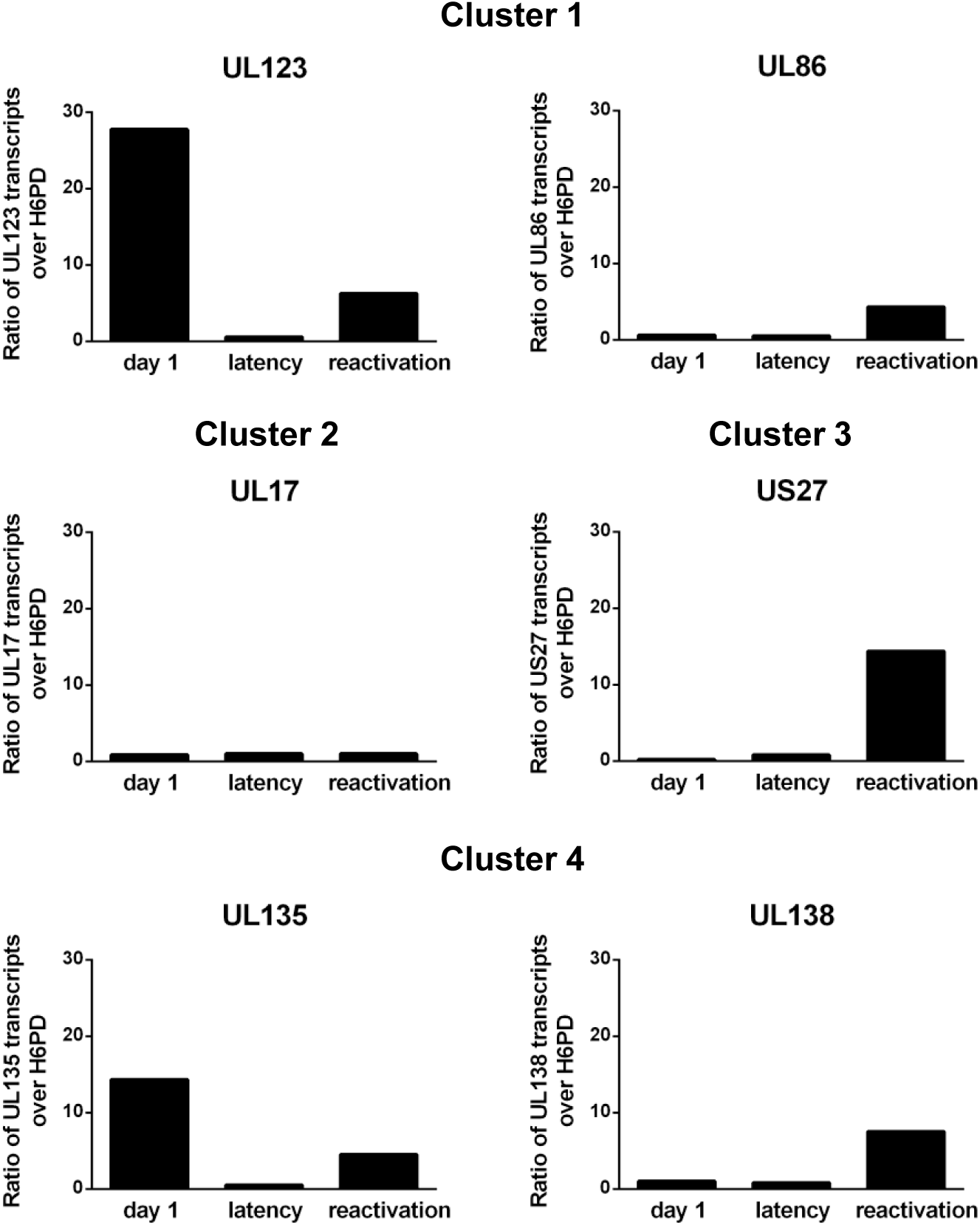
RT-qPCR confirmation of select transcripts in WT infection. THP-1 cells were infected with WT HCMV (MOI = 2) and cultured in suspension cell dishes. Total RNA was collected at 1 dpi during the establishment of latency. At 5 dpi, cell cultures were divided and treated with TPA (reactivation) to trigger re-expression of viral genes or DMSO (latency) to maintain the latent infection. Total RNA was collected from suspension cells (latency) and from adherent cells (reactivation) at 7dpi. RT-qPCR was performed to quantify viral transcripts and confirm the patterns of viral gene expression observed in the RNA-Seq analysis. Viral transcripts from each gene expression cluster (this study) and three canonical kinetic gene classes (IE, E, L) were selected. Data are expressed as ratio of viral transcripts over the cellular transcript H6PD and represent a single biological replicate analyzed in triplicate.

**Supplementary Figure 2.**
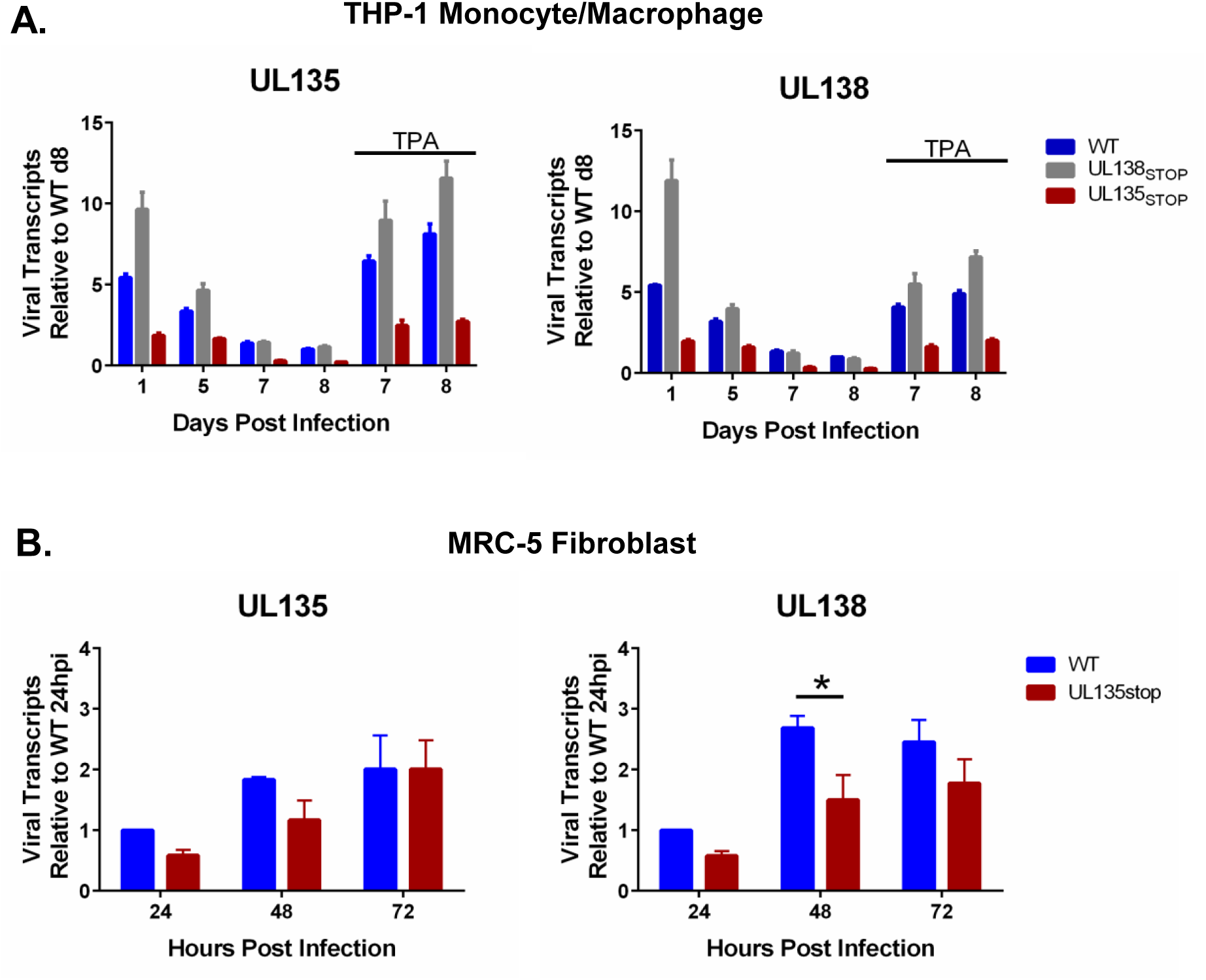
Comparison of representative UL*b*’ transcripts in hematopoietic versus replication-permissive cells. **A)** THP-1 cells were infected with WT, Δ*UL138*_STOP_, or Δ*UL135*_STOP_ HCMV (MOI = 2) and cultured in suspension cell dishes to establish latent infection. At 5 dpi, cells were treated with TPA to trigger re-expression of viral genes or with DMSO to maintain latent infection. Total RNA was isolated at the indicated time points and RT-qPCR was used to quantitate representative UL*b*’ transcripts *UL135* and *UL138*. Data are shown as the ratio of each viral transcript over cellular H6PD and represent a single biological replicate analyzed in triplicate and used to confirm viral gene expression patterns observed in the RNA-Seq analysis. **B)** MRC-5 fibroblasts were infected with WT or Δ*UL135*_STOP_ HCMV (MOI = 1) to establish replicative infection. Total RNA was collected at the indicated time points and RT-qPCR was used to quantify *UL135* and *UL138* transcripts. Data are expressed as fold change in viral transcripts over WT infection at 24 hours post infection (hpi). Error bars represent the SEM between three biological replicates analyzed in duplicate. Multiple t-tests (one per time point) were performed using the Holm-Sidak correction for multiple comparisons. Statistical significance where *, P < 0.05.

**Supplementary Data Set 1. Simple Enrichment Analysis.** Simple Enrichment Analysis (SEA) (*66*) was performed against the CIS-BP database of transcription factors (*67*) to identify transcription factor binding motifs that are significantly enriched in viral gene expression cluster 4 when compared to other viral gene expression clusters. The degree of significance for which each transcription factor is enriched in cluster 4 is expressed as a p-value. The percentage of cluster 4 HCMV genes associated with each predicted transcription factor binding site is shown (% HCMV c4 genes) as well as percent of HCMV genes from clusters 1, 2, and 3 (% HCMV c1, 2, 3, genes). Enrichment ratio represents the relative enrichment of each transcription factor binding site in cluster 4 versus clusters 1, 2, and 3. Figure quality images depicting the motif consensus sequences identified in cluster 4 genes were created using the WebLogo web-based application (*71, 72*).

**Supplementary Data Set 2. Differential Expression Analysis.** Transcription factors corresponding to the significantly enriched transcription factor binding sites in cluster 4 were ranked by degree of differential expression at each time point dependent on the presence of pUL135 in our RNA-Seq analysis. Negative binomial modeling of gene expression with the *DESeq2* package from R (*68*) was used to determine differential expression. Log fold change (logFC) of gene expression for *UL135*_STOP_ virus over viruses expressing the UL135 protein (WT and *UL138*_STOP_ averaged) is shown. Statistical significance where *, P < 0.05; **, P < 0.005 and ***, P < 0.0005.

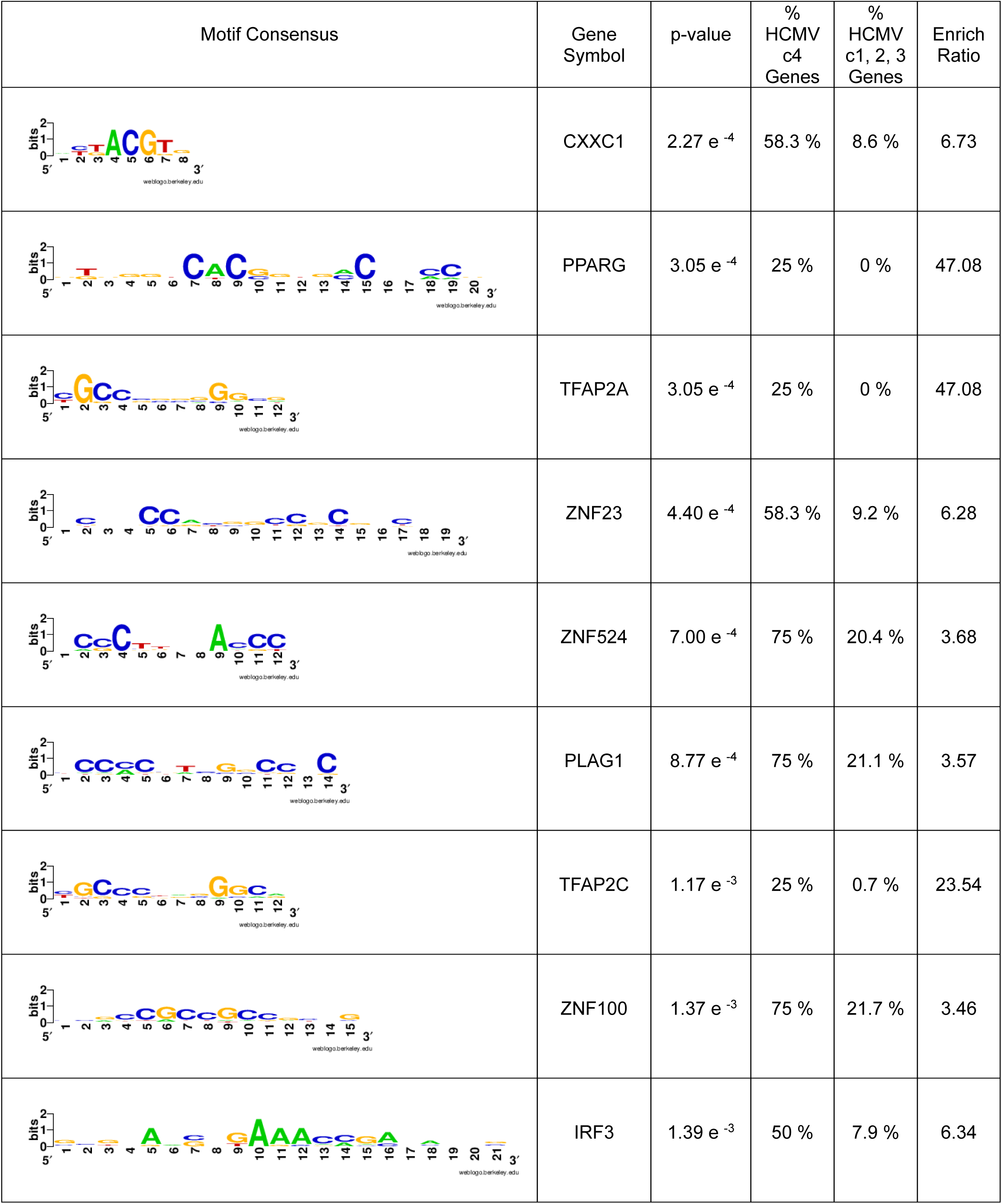
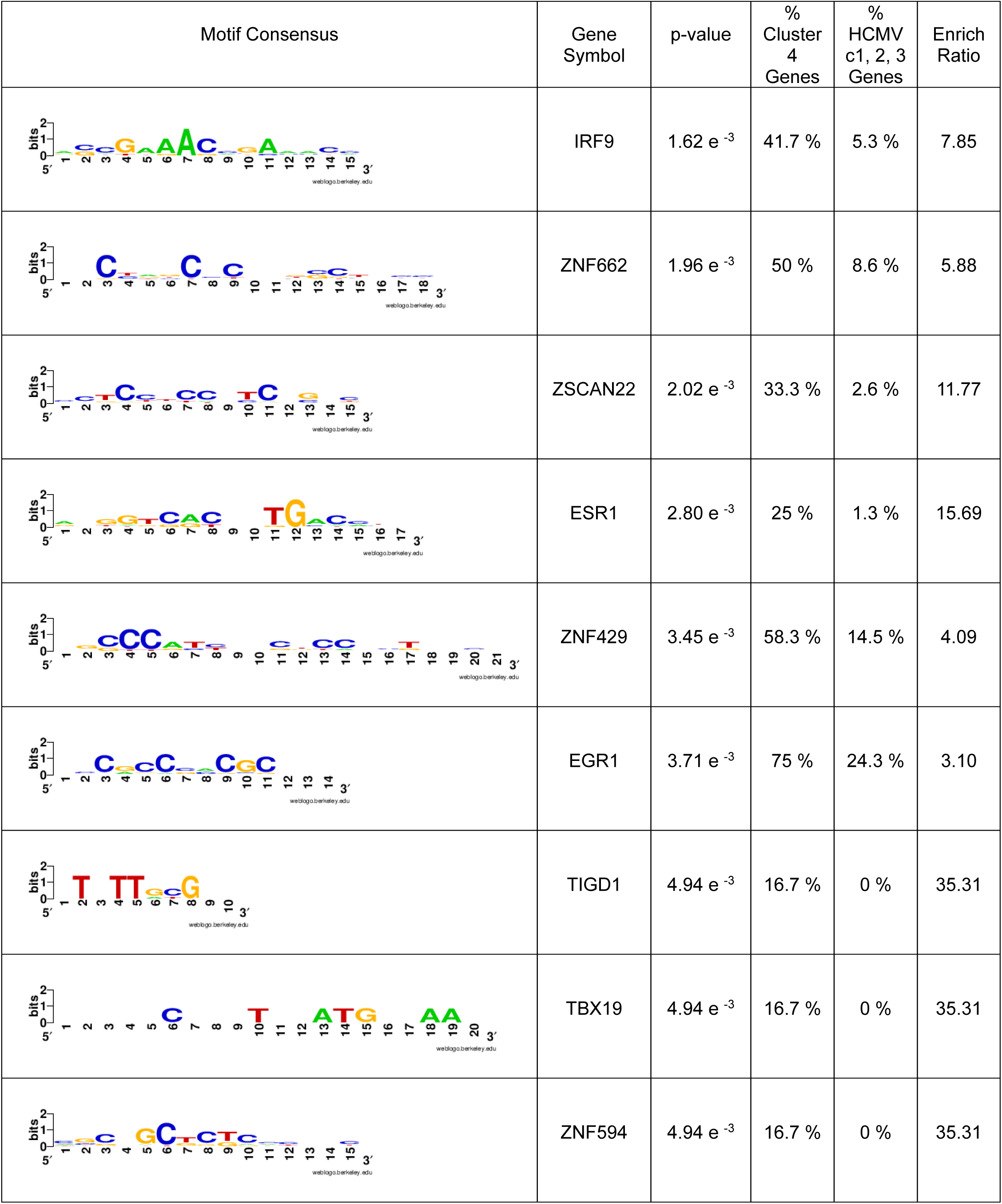
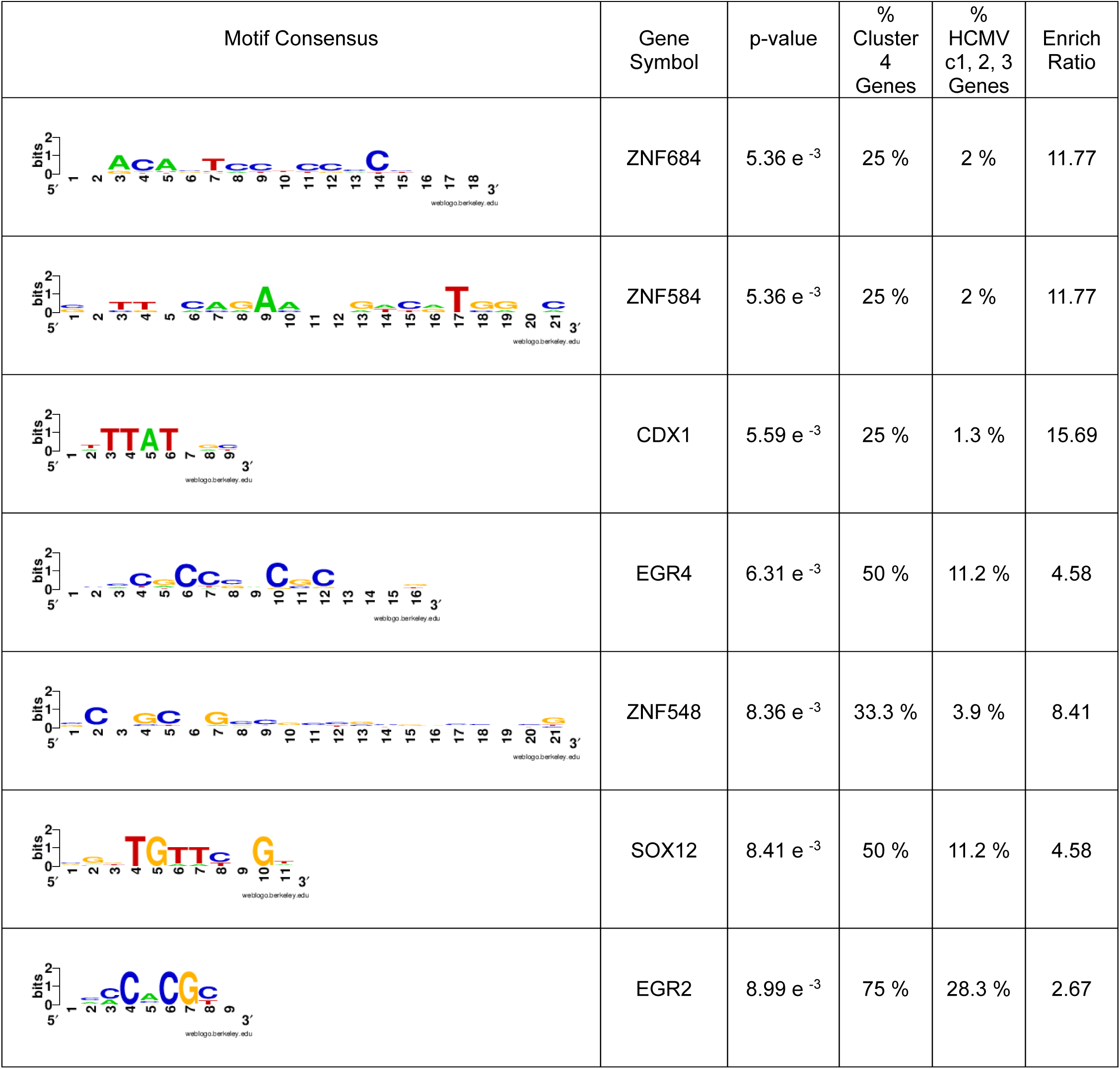

|  | gene symbol | ensembl | logFC UL135<sub>STOP</sub>/avg(WT+UL138<sub>STOP</sub>) | AvgExpr | t | p Value | adj p Value | B |
| --- | --- | --- | --- | --- | --- | --- | --- | --- |
| Day 1 undifferentiated | TFAP2C | ENSG00000087510 | -1.160202415 | 0.190043 | -4.15709 | 6.1343708 | 0.0008556 | 0.963543** |
|  | PPARG | ENSG00000132170 | 0.370753256 | 5.218841 | 3.523475 | 0.000607 | 0.0062923 | -1.20051* |
|  | CXXC1 | ENSG00000154832 | 0.157189768 | 5.641158 | 1.239288 | 0.217696 | 0.49348942 | -6.34767 |
|  | ZNF524 | ENSG00000171443 | 0.295048003 | 3.084672 | 0.908604 | 0.36541 | 0.64959332 | -6.69994 |
|  | ZNF100 | ENSG00000197020 | 0.088718788 | 4.539584 | 0.52154 | 0.602968 | 0.81698032 | -6.97596 |
|  | PLAG1 | ENSG00000181690 | 0.135495698 | 1.077953 | 0.49638 | 0.620549 | 0.8270811 | -6.98876 |
|  | IRF3 | ENSG00000126456 | -0.068330399 | 5.866038 | -0.43323 | 0.665636 | 0.85339777 | -7.01811 |
|  | ZNF23 | ENSG00000167377 | -0.065414956 | -2.07581 | -0.14147 | 0.887738 | 0.95776564 | -7.10203 |
|  | TFAP2A | ENSG00000137203 | -0.023972528 | 1.000728 | -0.12839 | 0.898057 | 0.96195426 | -7.1038 |
| Day 3 undifferentiated | TFAP2C | ENSG00000087510 | -0.798834005 | 0.190043 | -2.86228 | 0.004977 | 0.12550995 | -2.57596 |
|  | IRF3 | ENSG00000126456 | -0.198747266 | 5.866038 | -1.26011 | 0.210115 | 0.77750388 | -5.69694 |
|  | PPARG | ENSG00000132170 | 0.093190801 | 5.218841 | 0.885644 | 0.377609 | 0.88944352 | -6.08861 |
|  | CXXC1 | ENSG00000154832 | -0.099681904 | 5.641158 | -0.78589 | 0.433503 | 0.91307676 | -6.17022 |
|  | TFAP2A | ENSG00000137203 | 0.115498875 | 1.000728 | 0.618592 | 0.537377 | 0.94725838 | -6.28544 |
|  | ZNF524 | ENSG00000171443 | -0.197595561 | 3.084672 | -0.6085 | 0.544026 | 0.94947014 | -6.29152 |
|  | PLAG1 | ENSG00000181690 | -0.152047787 | 1.077953 | -0.55702 | 0.57857 | 0.95464582 | -6.32098 |
|  | ZNF100 | ENSG00000197020 | 0.081226818 | 4.539584 | 0.477498 | 0.63389 | 0.96730839 | -6.3614 |
|  | ZNF23 | ENSG00000167377 | 0.216359789 | -2.07581 | 0.467919 | 0.640705 | 0.96850953 | -6.36585 |
| Day 5 undifferentiated | CXXC1 | ENSG00000154832 | -0.225505293 | 5.641158 | -1.77789 | 0.077996 | 0.35376617 | -4.63076 |
|  | PPARG | ENSG00000132170 | 0.16329105 | 5.218841 | 1.551846 | 0.123375 | 0.43551737 | -4.98393 |
|  | PLAG1 | ENSG00000181690 | 0.411142146 | 1.077953 | 1.506194 | 0.134688 | 0.45417076 | -5.04969 |
|  | TFAP2A | ENSG00000137203 | -0.247783589 | 1.000728 | -1.32709 | 0.187041 | 0.52213634 | -5.28939 |
|  | ZNF524 | ENSG00000171443 | -0.387980027 | 3.084672 | -1.19479 | 0.234564 | 0.57186158 | -5.44757 |
|  | IRF3 | ENSG00000126456 | -0.148654527 | 5.866038 | -0.94251 | 0.347857 | 0.67176677 | -5.70418 |
|  | TFAP2C | ENSG00000087510 | -0.152089103 | 0.190043 | -0.54495 | 0.586818 | 0.82580627 | -5.98702 |
|  | ZNF23 | ENSG00000167377 | 0.229986979 | -2.07581 | 0.49739 | 0.619839 | 0.84332314 | -6.0108 |
|  | ZNF100 | ENSG00000197020 | 0.029735345 | 4.539584 | 0.174801 | 0.861535 | 0.95074931 | -6.11493 |
| Day 5.5 undifferentiated | PPARG | ENSG00000132170 | 0.182528313 | 5.218841 | 1.734669 | 0.085409 | 0.60943602 | -4.53355 |
|  | ZNF23 | ENSG00000167377 | -0.61479136 | -2.07581 | -1.3296 | 0.186212 | 0.76088461 | -5.10337 |
|  | TFAP2C | ENSG00000087510 | -0.365373922 | 0.190043 | -1.30916 | 0.193021 | 0.76862837 | -5.12827 |
|  | CXXC1 | ENSG00000154832 | -0.157928606 | 5.641158 | -1.24511 | 0.215555 | 0.79077285 | -5.20389 |
|  | PLAG1 | ENSG00000181690 | 0.196650438 | 1.077953 | 0.720417 | 0.472692 | 0.91338257 | -5.6831 |
|  | ZNF100 | ENSG00000197020 | 0.098327204 | 4.539584 | 0.578023 | 0.564349 | 0.93868392 | -5.76948 |
|  | TFAP2A | ENSG00000137203 | -0.104598384 | 1.000728 | -0.56021 | 0.576397 | 0.94173547 | -5.77897 |
|  | IRF3 | ENSG00000126456 | -0.041989988 | 5.866038 | -0.26623 | 0.790528 | 0.98121094 | -5.89272 |
|  | ZNF524 | ENSG00000171443 | -0.065643849 | 3.084672 | -0.20215 | 0.840147 | 0.98379825 | -5.90679 |
| Day 6 undifferentiated | TFAP2A | ENSG00000137203 | 0.20669957 | 1.000728 | 1.107047 | 0.270524 | 0.92464593 | -5.18008 |
|  | TFAP2C | ENSG00000087510 | 0.235149384 | 0.190043 | 0.842558 | 0.401179 | 0.95999793 | -5.4112 |
|  | ZNF524 | ENSG00000171443 | 0.183991866 | 3.084672 | 0.566605 | 0.572058 | 0.9773683 | -5.58622 |
|  | PPARG | ENSG00000132170 | 0.053351166 | 5.218841 | 0.507026 | 0.613082 | 0.98007183 | -5.61507 |
|  | ZNF23 | ENSG00000167377 | -0.183838918 | -2.07581 | -0.39759 | 0.691653 | 0.98577356 | -5.65976 |
|  | PLAG1 | ENSG00000181690 | 0.058973258 | 1.077953 | 0.216045 | 0.829326 | 0.99296086 | -5.71009 |
|  | ZNF100 | ENSG00000197020 | -0.035209278 | 4.539584 | -0.20698 | 0.836382 | 0.9940531 | -5.71182 |
|  | IRF3 | ENSG00000126456 | 0.028346522 | 5.866038 | 0.179724 | 0.857677 | 0.9977019 | -5.71659 |
|  | CXXC1 | ENSG00000154832 | 0.005258322 | 5.641158 | 0.041457 | 0.967002 | 0.99883043 | -5.73042 |
| Day 8 undifferentiated | CXXC1 | ENSG00000154832 | 0.284361045 | 5.641158 | 2.190365 | 0.030463 | 0.46870048 | -3.61542 |
|  | TFAP2A | ENSG00000137203 | 0.293741114 | 1.000728 | 1.537056 | 0.126955 | 0.62803435 | -4.68428 |
|  | TFAP2C | ENSG00000087510 | 0.418787284 | 0.190043 | 1.466046 | 0.145294 | 0.64438295 | -4.77901 |
|  | ZNF23 | ENSG00000167377 | 0.463869989 | -2.07581 | 0.980142 | 0.32902 | 0.77719166 | -5.31046 |
|  | ZNF524 | ENSG00000171443 | 0.306699358 | 3.084672 | 0.922769 | 0.358009 | 0.79291867 | -5.35956 |
|  | IRF3 | ENSG00000126456 | 0.127334203 | 5.866038 | 0.78877 | 0.431827 | 0.83183543 | -5.46288 |
|  | ZNF100 | ENSG00000197020 | -0.09500345 | 4.539584 | -0.54564 | 0.58634 | 0.89476917 | -5.60941 |
|  | PPARG | ENSG00000132170 | -0.05424017 | 5.218841 | -0.50362 | 0.615465 | 0.90637544 | -5.62936 |
|  | PLAG1 | ENSG00000181690 | -0.020320356 | 1.077953 | -0.07273 | 0.942143 | 0.98848408 | -5.74187 |
| Day 5.5 differentiated | TFAP2C | ENSG00000087510 | 0.464213495 | 0.190043 | 1.663311 | 0.098902 | 0.71881807 | -4.78323 |
|  | PLAG1 | ENSG00000181690 | -0.392897379 | 1.077953 | -1.43936 | 0.152696 | 0.85336045 | -5.10905 |
|  | TFAP2A | ENSG00000137203 | -0.09065915 | 1.000728 | -0.48555 | 0.628183 | 0.99991207 | -5.97955 |
|  | PPARG | ENSG00000132170 | 0.044178342 | 5.218841 | 0.419851 | 0.675357 | 0.99991207 | -6.00799 |
|  | IRF3 | ENSG00000126456 | -0.054938401 | 5.866038 | -0.34832 | 0.728217 | 0.99991207 | -6.03427 |
|  | ZNF524 | ENSG00000171443 | -0.110408863 | 3.084672 | -0.34001 | 0.734457 | 0.99991207 | -6.03701 |
|  | ZNF23 | ENSG00000167377 | -0.139931741 | -2.07581 | -0.30263 | 0.762705 | 0.99991207 | -6.04851 |
|  | CXXC1 | ENSG00000154832 | 0.027335438 | 5.641158 | 0.215513 | 0.829739 | 0.99991207 | -6.07011 |
|  | ZNF100 | ENSG00000197020 | -0.018806493 | 4.539584 | -0.11056 | 0.912157 | 0.99991207 | -6.0865 |
| Day 6 differentiated | PPARG | ENSG00000132170 | -0.308388064 | 5.218841 | -2.93078 | 0.00406 | 0.08845639 | -2.20072 |
|  | TFAP2A | ENSG00000137203 | 0.26509314 | 1.000728 | 1.419793 | 0.158303 | 0.53608525 | -5.24294 |
|  | TFAP2C | ENSG00000087510 | 0.345174731 | 0.190043 | 1.236787 | 0.21862 | 0.59882606 | -5.47478 |
|  | ZNF23 | ENSG00000167377 | -0.555554661 | -2.07581 | -1.20149 | 0.231965 | 0.60874713 | -5.51592 |
|  | PLAG1 | ENSG00000181690 | -0.295316164 | 1.077953 | -1.08187 | 0.281514 | 0.64704137 | -5.6467 |
|  | CXXC1 | ENSG00000154832 | 0.116318301 | 5.641158 | 0.917057 | 0.360982 | 0.70548616 | -5.80486 |
|  | IRF3 | ENSG00000126456 | 0.143897732 | 5.866038 | 0.912349 | 0.363444 | 0.70687041 | -5.809 |
|  | ZNF524 | ENSG00000171443 | 0.248932154 | 3.084672 | 0.76659 | 0.444855 | 0.75501121 | -5.92678 |
|  | ZNF100 | ENSG00000197020 | 0.084999029 | 4.539584 | 0.499673 | 0.618235 | 0.84770069 | -6.08987 |
| Day 8 differentiated | PPARG | ENSG00000132170 | -0.533937604 | 5.218841 | -5.07431 | 1.4632583 | 0.00027186 | 5.018128*** |
|  | TFAP2A | ENSG00000137203 | 0.374334215 | 1.000728 | 2.004869 | 0.047264 | 0.3996679 | -4.36901 |
|  | TFAP2C | ENSG00000087510 | 0.458430353 | 0.190043 | 1.64259 | 0.103128 | 0.56369915 | -4.99464 |
|  | ZNF100 | ENSG00000197020 | -0.231246487 | 4.539584 | -1.3594 | 0.176611 | 0.68461188 | -5.40089 |
|  | PLAG1 | ENSG00000181690 | -0.369958488 | 1.077953 | -1.35532 | 0.177902 | 0.68711741 | -5.4062 |
|  | ZNF23 | ENSG00000167377 | -0.610993475 | -2.07581 | -1.32139 | 0.188926 | 0.70162407 | -5.44978 |
|  | CXXC1 | ENSG00000154832 | 0.133474273 | 5.641158 | 1.052315 | 0.294804 | 0.78697692 | -5.75724 |
|  | IRF3 | ENSG00000126456 | 0.114885831 | 5.866038 | 0.728406 | 0.467807 | 0.85997536 | -6.03635 |
|  | ZNF524 | ENSG00000171443 | 0.200066211 | 3.084672 | 0.616106 | 0.53901 | 0.88553754 | -6.10965 |

